# A clubroot pathogen PBS3-like effector manipulates hormonal crosstalk to alter root morphology during colonization

**DOI:** 10.64898/2026.03.10.710858

**Authors:** Melaine Gonzalez-Garcia, Jiaxu Wu, Marina Silvestre Vañó, Soham Mukhopadhyay, Elisa Fantino, Robert Malinowski, Karolina Stefanowicz, Ian Major, Edel Pérez-Lopez

## Abstract

Salicylic acid (SA) and auxin are key regulators of plant immunity and development. The clubroot pathogen *Plasmodiophora brassicae* encodes *Pb*GH_3_, an effector related to the GH_3_ family involved in phytohormone homeostasis. Although *Pb*GH_3_ was proposed to conjugate auxin i*n vitro*, its biological function *in planta* has remained unclear. This study aimed to determine the *in vivo* role of *Pb*GH_3_ during host colonization.
We generated *Arabidopsis thaliana* and *Brassica napus* lines overexpressing *Pb*GH_3_ and characterized their developmental phenotypes, hormone profiles, gene expression, and susceptibility to infection. Structural modeling was performed to assess *Pb*GH_3_ similarity to plant GH_3_ proteins, and functional complementation was tested using the Arabidopsis *gh3*.*12* mutant.
The expression of *Pb*GH_3_ in Arabidopsis induced auxin-related developmental phenotypes without detectable accumulation of auxin conjugates. Instead, *Pb*GH_3_ structurally and functionally resembled GH_3.12_/PBS3 inducing increased conjugated SA levels, reduced jasmonic acid, suppressed *PIN*_*2*_ expression, and increased root hair number and infection. *Pb*GH_3_ complemented SA-related defects in the *gh3*.*12* mutant.
*Pb*GH_3_ functions as a modulator of SA metabolism rather than an auxin-conjugating enzyme, likely competing with host GH_3.12_/PBS3 to constrain effective SA accumulation. This reveals a novel strategy by which *P. brassicae* disrupts SA–auxin homeostasis to promote host colonization and ensure disease development.

**PLAIN LANGUAGE SUMMARY:** This study shows that the clubroot pathogen uses a protein called *Pb*GH_3_ to modify the plant’s salicylic acid balance. This alters root traits and increases susceptibility to infection. Arabidopsis and canola plants engineered to produce *Pb*GH_3_ showed similar changes, revealing that the pathogen uses this protein to disrupt hormone regulation and create conditions that support its colonization.

## INTRODUCTION

Plant hormones are central regulators of plant growth, development, and defense against biotic and abiotic stresses (Verma et al., 2016; Waadt et al., 2022). Precise control of hormone abundance, bioactivity, and spatial distribution is essential for appropriate signal perception and downstream responses. This regulation is achieved through coordinated networks of biosynthetic, catabolic, and modifying enzymes that collectively maintain hormonal homeostasis (Westfall et al., 2013). Among these, the Gretchen Hagen 3 (GH_3_) family of acyl acid amido synthetases plays a pivotal role by conjugating phytohormones to amino acids, thereby modulating hormone activity, stability, and cellular availability (Holland et al., 2019).

In *Arabidopsis thaliana*, 19 GH_3_ proteins have been identified, although only eight have been functionally characterized to date (Staswick et al., 2005; Jez, 2022). Based on substrate specificity and biological function, plant GH_3_ proteins are classified into three major groups: (*i*) enzymes that conjugate and inactivate indole-3-acetic acid (IAA); (*ii*) jasmonyl-isoleucine synthetases responsible for the formation of the bioactive jasmonic acid (JA) conjugate JA-Ile; and (*iii*) a Brassicaceae-specific group that catalyzes the synthesis of isochorismate–glutamate, a direct precursor of salicylic acid (SA) (Okrent et al., 2009; Holland et al., 2019). In Arabidopsis, this SA-associated GH_3_ group comprises only two functionally characterized members, *At*GH_3.12_ /PBS3 and *At*GH_3.7_, both of which contribute to SA biosynthesis and immune signaling (Westfall et al., 2013; Holland et al., 2019; Jez, 2022).

Beyond their established roles in development, GH_3_ proteins have emerged as key integrators of hormone-mediated immune responses during plant–pathogen interactions (Hui et al., 2019; Jez, 2022; Wojtaczka et al., 2022). For example, *At*GH_3.5_ functions as a regulatory hub linking SA- and auxin-dependent signaling during *Pseudomonas syringae* infection, thereby influencing resistance and susceptibility outcomes, respectively (Zhang et al., 2007). Similarly, *At*GH_3.12_/PBS3 plays a central role in SA biosynthesis by catalyzing the formation of the isochorismate–glutamate intermediate, a critical step in pathogen-induced SA accumulation (Chen et al., 2019). In rice, several GH_3_ proteins enhance resistance to *Magnaporthe grisea* and *Xanthomonas oryzae* by repressing auxin signaling pathways that otherwise favor pathogen colonization (Fu et al., 2011). Collectively, these studies highlight GH_3_ enzymes as crucial modulators of hormone crosstalk during immune responses.

In 2015, a GH_3_-like gene was identified in the genome of *Plasmodiophora brassicae*, an obligate biotrophic protist and the causal agent of clubroot disease in Brassica crops (Schwelm et al., 2015). This gene encodes a protein termed *Pb*GH_3_, which displays structural similarity to plant GH_3_ proteins, including GH_3.12_/PBS3, and exhibits enzymatic activity *in vitro*. However, despite these similarities, neither the original characterization (Schwelm et al., 2015) nor subsequent studies assessing *Pb*GH_3_ activity *in planta* (Smolko et al., 2024), directly tested whether this pathogen-derived GH_3_-like protein contributes to SA biosynthesis or interferes with host hormone metabolism acting as a GH_3.12_/PBS3-like effector.

It is well-known that active manipulation of host hormone metabolism by *P. brassicae* is a major determinant of clubroot progression, with perturbations in auxin and SA pathways playing central roles in host susceptibility. This raises the hypothesis that *Pb*GH_3_ functions as an effector protein that alters host hormone homeostasis to facilitate early steps of pathogen colonization. Here, we investigated the biological activity of *Pb*GH_3_ *in planta*. To this end, we generated *Arabidopsis thaliana* and *Brassica napus* transgenic lines constitutively expressing *Pb*GH_3_, following an experimental strategy previously employed to characterize the *P. brassicae* methyltransferase *Pb*BSMT (Bulman et al., 2019; Djavaheri et al., 2019). We assessed developmental and morphological phenotypes throughout plant growth and performed comprehensive hormonal profiling, including IAA, cytokinin (CK), abscisic acid (ABA), jasmonic acid, and salicylic acid, in roots overexpressiong *Pb*GH_3_ and wild-type plants. In parallel, we monitored *Pb*GH_3_ expression dynamics during clubroot infection and evaluated disease development following inoculation with the Canadian pathotype 3A. To further validate our findings, we complemented the *gh3*.*12/pbs3* knockout line with *Pb*GH_3_, phenotyped and determined SA concentration in the resulting lines to determine whether *Pb*GH_3_ has a similar functional role in SA metabolism.

## MATERIALS AND METHODS

### Plant material

*Arabidopsis thaliana* ecotype Columbia (Col-0) was used as the wild-type (WT) control. T-DNA insertion lines SALK_018225C (*pbs3/gh3*.*12*), SALK_094646C (*vas2/gh3*.*17*), SALK_033434C (*wes1/gh3*.*5*), and SALK_079153 (*gh3*.*15*) were obtained from the *Arabidopsis* Biological Resource Center (ABRC). All mutants were confirmed for homozygosity by PCR using primers designed with the SIGnAL T-DNA Primer Design tool (http://signal.salk.edu/tdnaprimers.2.html) (**Table S1**).

Plants were grown in controlled conditions of long-day photoperiod (16 h light/8 h dark, 21°C, 150 μmol/m^2^). For phenotyping and infection assays, plants were cultivated in a 3:1 (v/v) mixture of Veranda soil and vermiculite. For root analyses and hormone profiling, seeds were surface sterilized (70% ethanol with 0.005% Triton X-100, 3 min; 96% ethanol, 1 min), stratified for 3 days at 4°C in the dark, and sown on Murashige and Skoog (MS) medium agar plates for 5, 10 and 20 days as indicated for each experiment. For microscopy experiment, *A. thaliana* plants (Col-0 and transgenic Col-0:: 35S*Pb*GH_3_ lines) were grown under short-day conditions (9 h light /15 h dark) at 22 °C during the light period and 20 °C during the dark period, with a photosynthetically active radiation (PAR) of 120 µmol m^−2^ s^−1^ measured at canopy level.

Transgenic and wild-type *Brassica napus* L. cv. Westar seeds were surface-sterilized with ethanol 70% and germinated on autoclaved vermiculite for 7–10 days under controlled conditions (12 h light /12 h dark cycle, 24°C (day) /16°C (night), and 60% relative humidity). After germination, seedlings were transferred to a 32-cell seedling tray containing Veranda soil and grown in a greenhouse under long-day conditions for two days. Approximately one week after plants were transplanted into individual seedling pots and maintained in the greenhouse under the same conditions (Salih *et al*., 2023).

### *Plasmodiophora brassicae* inoculation and phenotyping

The *P. brassicae* 3A isolate was used in this study for infection assays (Javed et al., 2024). The canola cultivar Westar was used for pathogen propagation, and galls were stored at −20°C for resting spore extraction as previously described (Salih *et al*., 2024). Ten-day-old *Arabidopsis* plants were inoculated by pipetting 2 mL of a spore suspension (1 × 10^8^ spores mL^−1^) onto the root zone, while control plants received the same volume of distilled water (mock-inoculated). Fresh inoculum was prepared for each experiment. Clubroot phenotyping was conducted at 21 days post-inoculation (dpi) using a recently described scale that captures a more comprehensive range of host responses to clubroot (Gonzalez-Garcia et al., 2025).

For pathogen DNA quantification, roots from five *Arabidopsis* plants were pooled at 5-, 10-, and 21 dpi. DNA was extracted using a CTAB-based protocol (2% CTAB, 100 mM Tris–HCl, 20 mM EDTA, 1.4 M NaCl) (Doyle, 1990). Tissue was ground with one porcelain ball (2 × 40 s, 6 m s^−1^) and incubated at 65 °C for 1 h. DNA was precipitated using isopropanol and washed twice with 70% ethanol. The qPCR reactions (20 µL) contained 15 ng DNA, 0.25 µM primers, and Luna qPCR Master Mix (NEB, CAD) and were run on a CFX Opus system (BioRad, CAD). Host and pathogen targets housekeeping genes were AtSK11 (At5g26751) and Pb18S (ENSRNAT00050137123), respectively. The primers used for the quantitative analysis are described in Table S1. Relative pathogen DNA titer was calculated as ΔCt = Ct_{Pb18S}_ − Ct_{AtSK11}_. Three technical replicates were performed for each sample and were averaged.

Root colonization was evaluated at 3 dpi using ten-day-old seedlings infected with pathotype 3A. Roots were collected, rinsed twice with distilled water, and mounted on slides for microscopy. Images were captured with an Axio microscope (Zeiss, Germany) and analyzed in ImageJ (NIH). Twenty plants per line were evaluated.

### Generation of transgenic *A. thaliana* and *B. napus* lines

Gateway cloning was used to generate the expression construct. The coding sequence of the hypothetical protein PBTT_05886/PBRA_002260 (GenBank: CEP01995) was synthesized and cloned into the entry plasmid pENTR-Kozak (Twist Bioscience, USA), yielding pENTR-Kozak_*Pb*GH_3_ (kanamycin selection). The entry clone was double digested with SspI and ClaI, treated with phosphatase, and recombined into the binary destination vector pMDC32. Recombinant plasmids were introduced into *Escherichia coli* TOP10 by heat shock to conserve the construction. Plasmid DNA was purified with GeneJET Plasmid Miniprep Kit (Thermo Fisher Scientific, CAD) from *E*.*coli* TOP10 cells and electroporated into *Agrobacterium tumefaciens* GV3101. Transformed colonies were confirmed by colony PCR and used for plant transformation. The resulting construct (pMDC32_ *Pb*GH_3_ (**Figure S1**) was verified by whole-plasmid sequencing (Plasmidsaurus Inc., USA) and deposited and available in Addgene (plasmid #232335; RRID: Addgene_232335).

Approximately 5-week-old Col-0 and SALK_018225C plants were transformed with *A. tumefaciens* GV3101 harboring pMDC32_ *Pb*GH_3_ using the floral dip method (Clough & Bent, 1998). Plants were grown under long-day conditions in the greenhouse as described above. The seeds were collected and screened in MS medium supplemented with 50mg/L of hygromycin (VWR, CAD) until the obtention of the homozygous T_3_ generation, using T_4_ seeds in further experiments.

Transformation of *Brassica napus* L. cv. Westar was performed using a modified cotyledon explant protocol mediated by *A. tumefaciens* (Cardoza & Stewart, 2006; Bhalla & Singh, 2008; Zhang et al., 2020). *A. tumefaciens* strain GV3101 harboring the binary vectors pMDC32-*Pb*GH_3_ was cultured in LB medium supplemented with kanamycin (50 mg/L) and rifampicin (50 mg/L) for 48 h at 28 °C with shaking (200 rpm). Surface-sterilized seeds of *B. napus* were germinated on half-strength MS medium for 4 days in the dark, and cotyledonary explants were excised and immersed in *Agrobacterium* suspension (OD_600_ = 0.4-0.6) with 100mM Acetosyringone for 30 min before being transferred to co-cultivation medium. Explants were incubated for 2 days at 25 °C under dim light (~660 lux), followed by selection on callus induction medium containing 50 mg/L of hygromycin. Shoots regenerated after 4 to 5 weeks were transferred to shoot induction medium containing hygromycin (50 mg/L) for 2 to 4 weeks, and subsequently to shoot growth medium with 50 mg/L of hygromycin. Root induction was performed on half-strength MS medium supplemented with 10 g/L sucrose and 100 µL IBA stock (10 mg/mL). Rooted plantlets were acclimated in soil under mist conditions for 1 week and then transferred to greenhouse conditions. *Arabidopsis* and canola transgenic plants were analyzed to check positive transformants by PCR and RT-qPCR (**Table S2** and **Table S3**). Three biological replicates were done for each independent line and the primers used are listed in supporting files (**Table S1**).

### Morphological characterization of transgenic plants

Three independent lines for each *Arabidopsis* background (Col-0 or SALK_018225C) as well as the wild-type ecotype Col-0 and the mutant *gh3*.*12*/*pbs3* (SALK_018225C) were analyzed to check the presence of any distinctive trait related to the expression of the transgene. Plants were grown during the entire plant life cycle under controlled long-day conditions. Flowering time was expressed as the number of leaves at the bolting stage; leaves were photographed at that stage. At the maturity stage (~42–45 d), floral morphology, number of branches, plant height, and silique number were recorded using siliques 11–15 on the main inflorescence (numbered from the first silique). Five-day-old seedlings were analyzed for hypocotyl length and root hair density, and 10-day-old seedlings for rosette size, primary root length, and lateral root number and length. Each trait was measured in 20–30 plants per line.

In addition, transgenic *B. napus* L. cv. Westar lines overexpressing *Pb*GH_3_ were evaluated under long-day growth conditions. Three independent T_2_ lines were selected based on *Pb*GH_3_ expression. Morphological traits including plant height, number of branches, and time to flower were recorded. Root architecture parameters as primary root length, lateral root number, and lateral root length were analyzed in 10-day-old seedlings grown on MS agar plates; while the number of root hairs was analyzed in 5-day-old seedlings.

### Microscopy

Plant hypocotyls from Col-0 and Col-0::35S*Pb*GH_3_ lines were collected, thoroughly cleaned, and fixed in 4% (w/v) paraformaldehyde in 1× PBS as previously described (Fuchs *et al*., 2014). Fixed samples were embedded in 4% (w/v) agarose and sectioned using a vibratome to obtain transverse sections of 30-40 µm. Next, samples were cleared and stained with 5% (v/v) solution of Calcofluor white for 5 min in the dark. Cleared sections were mounted in clearing solution (Singh et al., 2022). The resulting sections were visualized using an epifluorescence microscope (Zeiss, USA), as previously described (Singh et al., 2022).

### Hormone content analysis

*Arabidopsis* plants (Col-0, SALK_018225C, transgenic lines: Col-0::35S*Pb*GH_3_ and SALK_018225C::35S*Pb*GH_3_) were grown vertically on MS medium for 20 days under long-day conditions in a growth chamber. For auxin and cytokinin (CK) analysis, roots of 40 plants per line were pooled, lyophilized, and ground. For SA and JA analysis, roots of 80 plants per line were pooled and ground in liquid N_2_. Three biological replicates were prepared per assay. Hormone quantification was performed at the National Research Council of Canada (Saskatoon, SK) using HPLC/UPLC–ESI–MS/MS on a Waters ACQUITY UPLC coupled to a Micromass Quattro Premier XE triple quadrupole mass spectrometer. Quantification was performed in MRM mode using isotope-labelled internal standards, and hormone levels were determined from analyte/internal standard peak area ratios relative to calibration curves. QC samples and blanks were included in each run. The method followed a modified protocol from Lulsdorf et al. (2013).

### Auxin treatment

Col-0 and Col-0::35S*Pb*GH_3_ lines were sown on MS agar supplemented with increasing concentrations (0.01–1 µM) of indole-3-acetic acid (IAA) (Sigma, CAD), 2,4-D (PhytoTech, CAD), or 1-naphthaleneacetic acid (NAA) (Sigma, CAD). After stratification, plates were grown vertically positioned in a growth chamber under long-day conditions (16h light, 8h dark), 21 degrees day and night. Ten-day-old seedlings were photographed, and primary root length was measured using ImageJ. Fifteen to twenty seedlings per line were analyzed.

### RNA isolation and RT-qPCR analysis

For RNA extraction, samples were collected and flash frozen in liquid N_2_. Tissue was grinded using mortar and petle, then RNA was extracted using the E.Z.N.A.® Plant RNA Kit (Omega, CAD) and quantified using Nanodrop OneC (Thermo Fisher Scientific, CAD). Residual genomic DNA was removed with TURBO DNase (Thermo Fisher Scientific, CAD), and first-strand cDNA was synthesized with SuperScript IV Reverse Transcriptase (Thermo Fisher Scientific, CAD).

RT-qPCR was performed on a CFX Opus Real-Time PCR System (Bio-Rad, CAD) in 20 µL reactions containing Luna Universal Probe qPCR Master Mix (NEB) and 0.2 µM primers. Non-template controls (NTCs) and no-reverse-transcriptase controls (NRTs) were included. Primer amplification efficiency was determined using standard curves, and all primer pairs showed efficiencies >90% (R^2^ > 0.99). Expression was normalized to the housekeeping genes previously mentioned. Data was analyzed with CFX Maestro Software 2.3 (Bio-Rad, CAD). Additionally, to quantify *Pb*GH_3_, *At*TAA1, *At*PIN1, and *At*PIN2 expression the housekeeping gene *At*TIP41 (*At4g34270)* was used. A similar procedure was followed to quantify *Pb*GH3 expression during disease progress in Arabidopsis at 0-, 2-, 6-, 10-, 12-, 16-, 18-, and 21 dpi.

### Structural analysis

Phylogeny and structural comparisons of *Pb*GH_3_ were performed using the telomere-to-telomere reference genome of *P. brassicae* pathotype 3A (Javed et al., 2024). Nineteen GH_3_ protein sequences from TAIR were downloaded (*AT*1g48690 was excluded as it encodes a truncated 190 aa fragment) and aligned with MAFFT (v7.525, --auto mode). Alignment columns with >20% gaps were trimmed, reducing the alignment from 731 to 566 columns. Phylogenetic trees were inferred with IQ-TREE2 (v2.0.7) using the LG+I+G4 substitution model, selected by BIC via ModelFinder, with 1000 ultrafast bootstrap replicates and 1000 SH-aLRT replicates, and visualized with iTOL. For structural comparisons, the AlphaFold2-generated model of PbGH3 was retrieved from UniProt and processed using FoldSeek to identify the most structurally similar Arabidopsis hits. The top Arabidopsis hits were further aligned to PbGH3 using TM-align and visualized in PyMOL..

### Statistical analyses

All statistical analyses were performed in R (R Core Team) within RStudio. Data were tested for normality and homoscedasticity before selecting statistical models. Parametric data were analyzed with two-tailed t-tests comparing transgenic lines to controls. Nonparametric data were analyzed using the non-parametric model Wilcoxon rank-sum test. Statistical significance was defined as *p* < 0.05.

## RESULTS

### *Pb*GH_3_ induces high-auxin phenotype in *Arabidopsis* and canola

To investigate the role of *Pb*GH_3_ in hormone homeostasis, we generated three independent *Arabidopsis* T4 lines expressing *Pb*GH_3_ under the control of the constitutive promoter 35S from cauliflower mosaic virus (CaMV) (**Figure S2, Table S2**). Col-0::35S*PbGH*_*3*_ lines, the control Col-0, and Col-0 expressing the empty vector (EV) were grown under normal long-day conditions. 10 dpg, over-expression of *Pb*GH_3_ led into plants with smaller, narrower, downward-bending leaves characteristic of epinasty (Sandalio et al., 2016), while the control EV was indistinguishable from the wild type Col-0 (**Figure 1A-B**). Petiole length of the leaves at positions 5 and 8 from the rosette were measured as well, but no differences were found compared to Col-0 (**Table S4**). Under long-day conditions, transgenic plants showed a different plant architecture supported by the appearance of bushier plants as a consequence of reduced apical dominance (**Figure 1C-D**). Plant height and lateral branch number were not significantly different for Lines 1.5 and 4.1, although the three lines followed a similar trend (**Figure 1F-G**). Regarding reproductive features, Col-0::35S*Pb*GH_3_ plants displayed early flowering, measured as the number of leaves at the bolting stage (**Figure 1E**); as well as greater number siliques with a reduced size. No obvious defects were observed on flower architecture, or floral organs (**Figure S3, Table S4**). Overexpression of *PbGH*_*3*_ in *B. napus* also led to high-auxin phenotypes as it was observed for *Arabidopsis* (**Figure 1H-I, Figure S2; Table S4**). We analyzed three independent canola T_2_ lines, expressing *PbGH*_*3*_ after transformation, and all of them showed a clear reduction in apical dominance together with an increase in lateral branching (**Figure 1H-I, Table S4**), consistent with the phenotype observed in *Arabidopsis* and suggesting that *Pb*GH_3_ regulates auxin-dependent developmental pathways that are conserved across Brassica crops.

**FIGURE 1:**
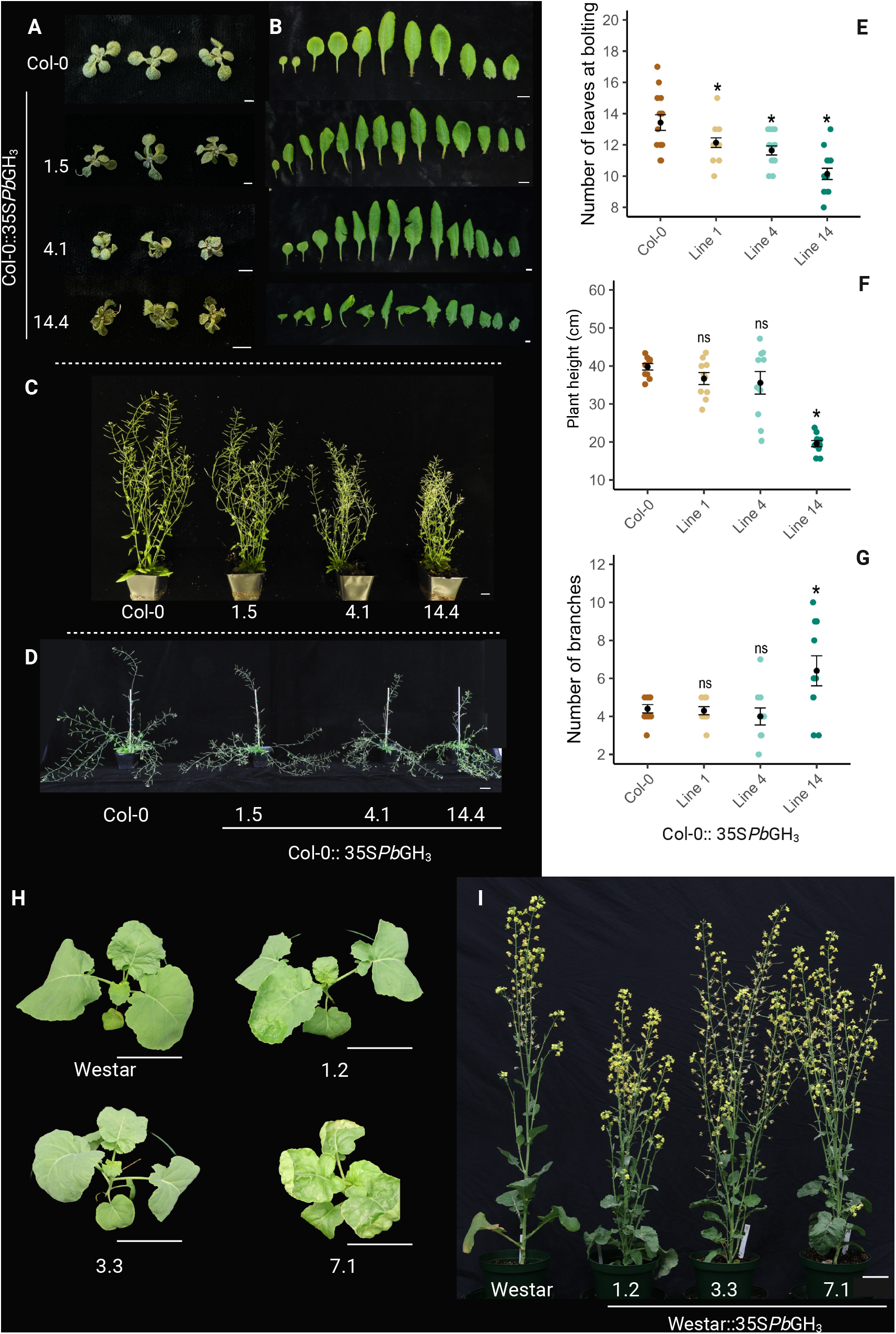
Morphological alterations induced by *Pb*GH_3_ overexpression in *Arabidopsis* and canola under long-day conditions. (**A**) Reduced rosette size in 10-day-old *Arabidopsis* seedlings (scale bar = 0.5 cm). (**B**) Epinastic leaves in *Arabidopsis* plants at the bolting stage (scale bar = 1 cm). (**C**) Representative *Arabidopsis* plants at flowering, illustrating overall plant height (scale bar = 1 cm). (**D**) Reduced apical dominance in *Arabidopsis* plants overexpressing *Pb*GH_3_ (scale bar = 1 cm). (**E**) Reduced number of rosette leaves at bolting stage in *Arabidopsis* plants (n = 10). (**F**) Plant height of *Arabidopsis* at flowering (n = 10). (**G**) Number of lateral branches in *Arabidopsis* plants (n = 10). (**H**) *B. napus* plants at 24 dpg (scale bar = 10 mm). (**I**) *B. napus* plants at 50 dpg (scale bar = 10 mm). Asterisks indicate significant differences relative to Col-0 (*p* < 0.05); ns, not significant (Student’s t-test for panels E and F; Wilcoxon test for panel G).

### *Pb*GH_3_ does not conjugate auxin in vivo

Previous *in vitro* studies demonstrated that *Pb*GH_3_ can conjugate auxin (Schwelm et al., 2015; Smolko et al., 2024). To further investigate if *Pb*GH_3_ mediates auxin conjugation *in planta, Arabidopsis* seedlings overexpressing *Pb*GH_3_ and WT Col-0 were grown on MS agar supplemented with increasing concentrations (0.01–1 µM) of the auxins IAA, NAA, or 2,4-D. Because exogenous auxin application has been proved to inhibit primary root growth in *Arabidopsis* (Ravelo-Ortega et al., 2021), we hypothesized that auxin conjugation by *Pb*GH_3_ would attenuate this inhibitory effect, resulting in reduced sensitivity to exogenous auxin and comparatively longer primary roots in transgenic lines (Fukui et al., 2022).

At 10 dpg, primary root growth inhibition in response to IAA was comparable between Col-0 and *Pb*GH_3_ transgenic lines (**Figure 2A, Table S5, Figure S4A**). Similarly, responses to NAA were indistinguishable among the overexpressing lines and the control across all tested concentrations (**Figure 2B, Table S5, Figure S4B**). In response to 2,4-D (Seifertová et al., 2014), overexpressing lines 4.1 and 14.4 exhibited a small increase in primary root length at the lowest concentration tested (0.01 µM) (**Figure 2C, Table S5**). Whereas no significant differences were observed at higher concentrations (**Figure 2C, Table S5, Figure S4C**).

**FIGURE 2:**
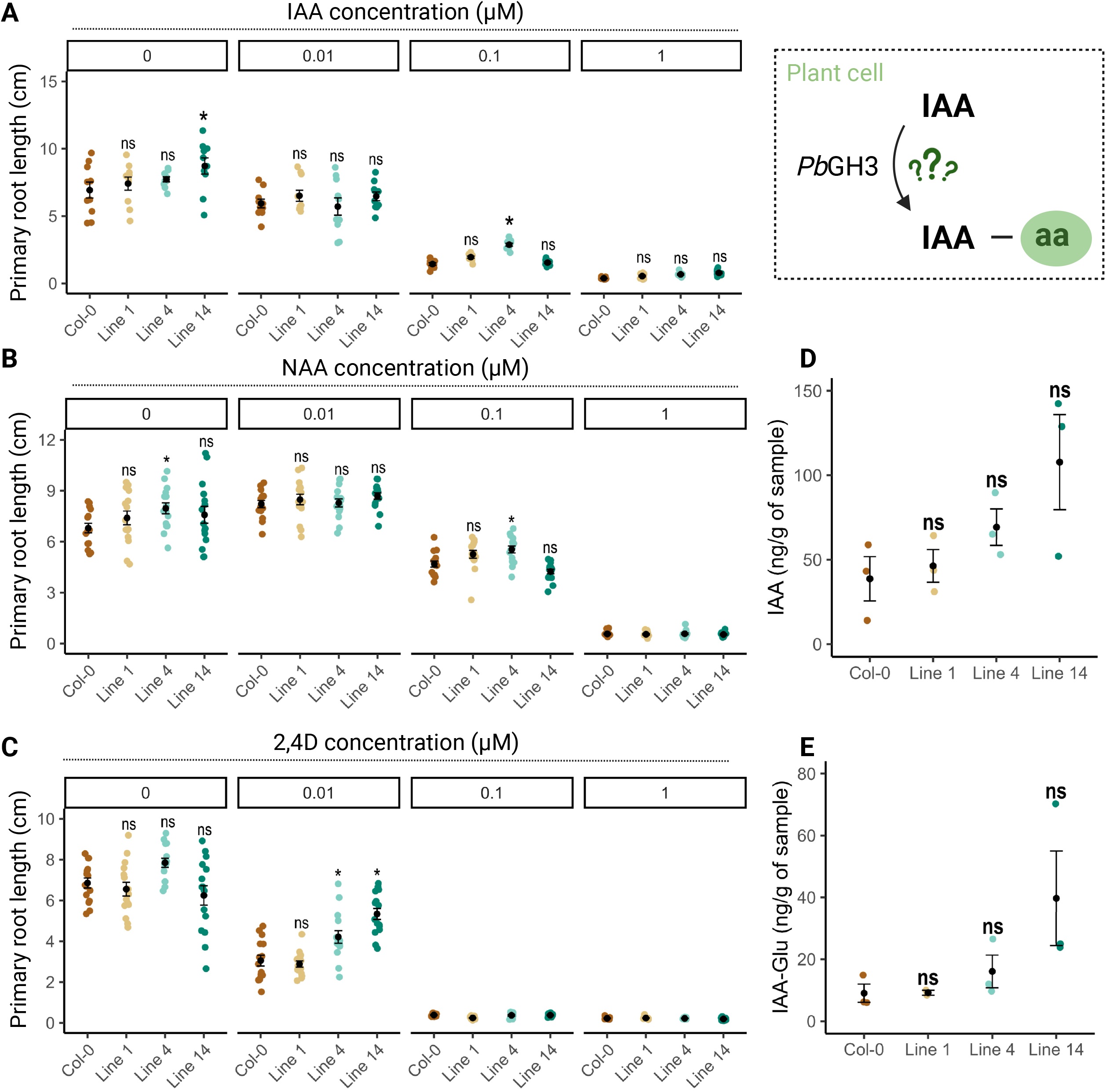
*Pb*GH_3_ overexpression does not affect auxin sensitivity or auxin conjugation *in planta*. Primary root length of wild-type Col-0 and Col-0::35S*Pb*GH_3_ *Arabidopsis* lines was measured at 10 days post-germination on MS medium supplemented with increasing concentrations (0–1 µM) of indole-3-acetic acid (C, IAA), naphthaleneacetic acid (D, NAA), or 2,4-dichlorophenoxyacetic acid (E, 2,4-D). Individual points represent single seedlings. Quantification of free IAA (A) and the conjugated form IAA–glutamate (B, IAA–Glu) was performed on roots of 20-day-old plants, revealing no significant differences between Col-0 and Col-0::35S*Pb*GH_3_ lines. Statistical significance was assessed relative to Col-0 within each treatment (P < 0.05; ns, not significant). Schematic representation of possible auxin conjugation by *Pb*GH_3_ is shown as reference in the top right.

To further assess the role of *Pb*GH_3_ in auxin conjugation *in planta*, hormone profiling was performed on roots of the three independent *Arabidopsis Pb*GH_3_ overexpressing lines and in the control Col-0 at 20 dpg. A higher level of free auxin was observed in Col-0::35S*Pb*GH_3_ lines compared with the control Col-0, being line 14.4 the one that showed the greater amount of free IAA. Although this trend was observed in all the overexpressing lines compared to the control, these differences were not statistically significant (**Figure 2D, Table S6**). Among auxin conjugates, only IAA-glutamate was detected in the *Pb*GH_3_ overexpressing lines compared to the control following the same trend showed for auxin (**Figure 2E, Table S6**). Notably, conjugated auxin levels were substantially lower than those of bioactive free IAA.

Other hormones were detected as well as CK and ABA. From the different forms of CK analyzed none of the bioactive free base CKs (Zeatin, Dihydrozeatin, Isopentenyladenine) were found in these samples. However, their biosynthetic precursor Zeatin riboside (ZR), both *cis-* and *trans-* isomers, and isopentenyladenine riboside (iPR), as well as the Z catabolism product cis-Zeatin-O-glucoside (ZOG) were detected (**Figure S5, Table S6**). These results were consistent with those for auxin, showing that overall cytokinin content was the greatest in line 14.4. Additionally, we detected the presence of ABA and some of its known catabolites in all samples, with 8⍰-hydroxylation (phaseic acid, dihydrophaseic acid) predominating over free ABA (**Figure S5, Table S6)**. Together with the lack of tolerance to exogenous auxin, these hormone profile suggest that *Pb*GH_3_ influences auxin homeostasis *in planta*, but not through direct auxin conjugation under the conditions tested.

### *Pb*GH_3_ induces changes in root phenotype in *Arabidopsis* and canola

Clubroot is recognized as a disease that primarily affects root tissues, with the clubroot pathogen initiating colonization in the root hairs (Javed *et al*., 2024). We assessed the impact of *PbGH*_*3*_ overexpression on root morphology by comparing three independent *Pb*GH_3_ transgenic lines in *Arabidopsis* and canola with their respective WT backgrounds, Col-0 and Westar. In *Arabidopsis*, primary root length measured at 10 dpg is not statistically significant between *Pb*GH_3_ transgenic lines and Col-0 (**Figure 3A, Table S7**). All *Pb*GH_3_ overexpressing *Arabidopsis* lines exhibited an increase in root hair density at 5 dpg (**Figure 3B–C, Table S7**). The analysis of lateral root development revealed that lines 1.5 and 14.4 produced fewer lateral roots but significantly longer lateral roots relative to the WT (**Figure 3D-E, Table S7**). Hypocotyl length was not affected by *Pb*GH_3_ overexpression in *Arabidopsis* (**Table S6**).

**FIGURE 3:**
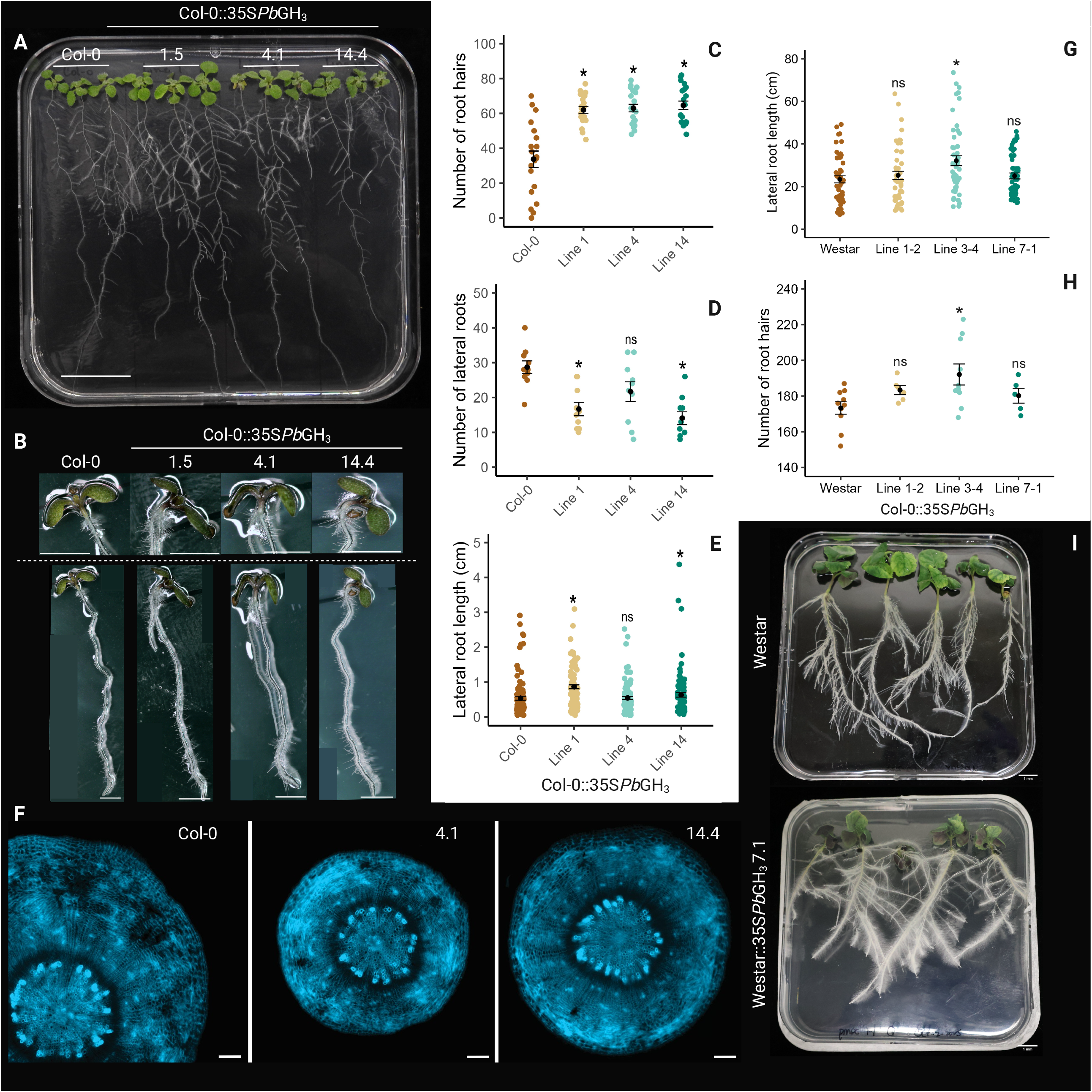
*Pb*GH_3_ overexpression alters root architecture without affecting vascular organization. (**A**) Root system architecture of Col-0 and Col-0::35S*Pb*GH_3_ seedlings grown vertically on MS medium at 10 dpg (scale bar = 1 cm). (**B**) Representative images of primary roots from wild-type Col-0 and independent Col-0::35S*Pb*GH_3_ lines (1.5, 4.1 and 14.4) at 10 dpg, highlighting increased root hair density in transgenic lines (scale bars = 0.5 cm). (**C**) Quantification of root hair number per primary root. (**D**) Number of lateral roots per seedling. (**E**) Average lateral root length at 10 dpg. In panels C–E, individual points represent single plants. Asterisks indicate significant differences relative to Col-0 (P < 0.05); ns, not significant (one-way ANOVA followed by post hoc test). (**F**) Transverse sections of primary roots stained with Calcofluor white showing root tissue organization in Col-0 and Col-0::35S*Pb*GH_3_ Lines 4.1 and 14.4. (scale bars = 100 µm). (**G**) Primary root length of *Brassica napus* (Westar) and three independent Westar::35S*Pb*GH_3_ transgenic lines (L1-2, L3-4 and L7-1). Individual data points represent single plants; black bars indicate mean ± SE. Asterisks indicate significant differences compared with Westar (Student’s t-test; P < 0.05), and ns indicates not significant. (**H**) Number of root hairs in *B. napus* (Westar) and Westar::35S*Pb*GH_3_ transgenic lines (L1-2, L3-4 and L7-1). Individual data points represent single roots; black bars indicate mean ± SE. Asterisks indicate significant differences compared with Westar (Student’s t-test; P < 0.05), and ns indicates not significant. (**I**) Representative images of root systems of *B. napus* Westar and Westar::35S*Pb*GH_3_ line L7-1 grown under long-day conditions. Additional images of independent Westar::35S*Pb*GH_3_ lines are shown in Figure S6.

Because clubroot infection is frequently associated with alterations in root vascular organization, including changes in xylem and phloem differentiation driven by pathogen-induced metabolic reprogramming (Malinowski et al., 2019; Silvestre Vañó et al., 2024), we next examined whether *Pb*GH_3_ expression affected vascular tissue organization. Transverse sections of hypocotyls stained with Calcofluor White revealed well-defined, radially organized structures in both wild-type Col-0 and Col-0::35S*Pb*GH_3_ lines 4.1 and 14.4, with clearly distinguishable cortical layers and a centrally positioned vascular cylinder (**Figure 3F**). No obvious differences in overall tissue organization or vascular patterning were detected between Col-0 and *Pb*GH_3_ over-expressing lines, indicating that *Pb*GH_3_ overexpression does not cause major alterations in root anatomical organization.

Similar altered phenotypes were observed in canola *Pb*GH_3_ over-expressing lines, where all lines displayed significant increase on lateral root length and root hair density compared to the Westar control (**Figure 3G–H, Figure S6, Table S7**). Being the line 3.4 the one that showed the greatest increase (**Figure 3G-I, Figure S6, Table S7**). Together, these results indicate that *Pb*GH_3_ overexpression modifies root architecture in both *Arabidopsis* and canola, potentially increasing root surface, by inducing root length and root hair development. These changes could influence early stages of pathogen interaction leading to greater root colonization.

### *Pb*GH_3_ overexpression does not affect disease progress but facilitates pathogen root colonization

As *Pb*GH_3_ over-expression produced consistent developmental phenotypes in both *Arabidopsis* and canola, we focused subsequent infection assays on Arabidopsis as a genetically tractable model for studying *Pb*GH_3_ function during *P. brassicae* infection. Given that *Pb*GH_3_ over-expression promotes root hair formation, a key entry site for *P. brassicae*, we next evaluated clubroot disease progression in Col-0::35S*Pb*GH_3_ transgenic lines in comparison with Col-0. Disease phenotyping at 21 dpi revealed that lines 1.5 and 4.1 exhibited similar level of clubroot susceptibility as the WT Col-0, whereas line 14.4 displayed reduced susceptibility, with a subset of plants remaining symptom-free (**Figure 4A–B, Figure S7, Table S8)**. Pathogen DNA quantification revealed comparable *P. brassicae* titers across all overexpressing lines in comparison to Col-0 at 21 dpi (**Figure 4C, Table S9**). Therefore, the altered disease phenotype observed in line 14.4 is more likely attributable to *Pb*GH_3_ overexpression-associated developmental changes rather than to differences in pathogen load (**Figure 4C, Table S9**).

**FIGURE 4:**
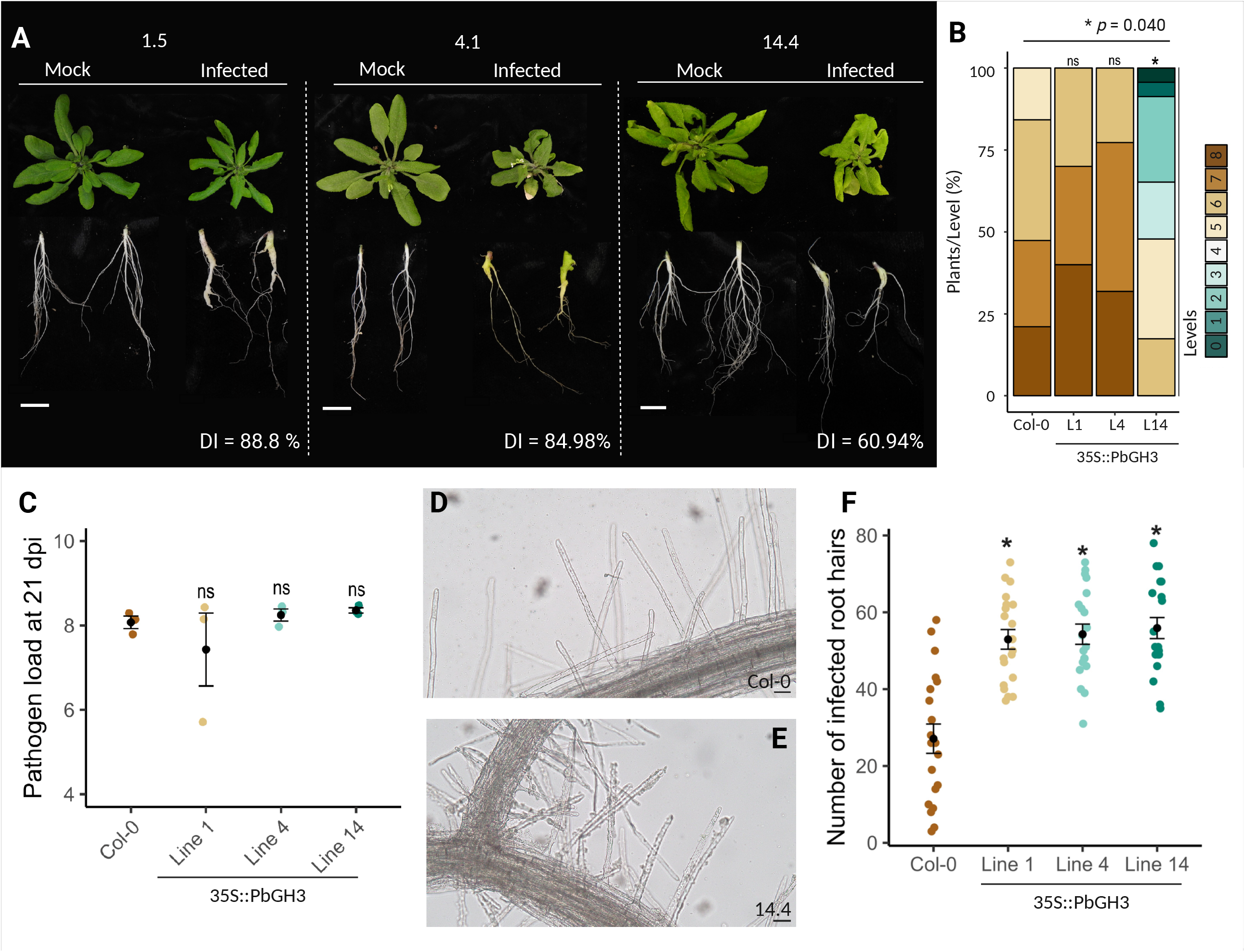
*Pb*GH_3_ overexpression promotes early root hair colonization but does not influence clubroot severity in *Arabidopsis*. (**A**) Representative images of Col-0::35S*Pb*GH_3_ *Arabidopsis* lines 1.5, 4.1 and 14.4 inoculated with water or Plasmodiophora brassicae pathotype 3A. Whole plants and corresponding root systems are shown at 21 dpi. Disease index (DI) values for each line are indicated (scale bars = 1 cm). Wild type col-0 is presented in Figure S7. (**B**) Distribution of disease severity levels at 21 dpi expressed as the percentage of plants per category. (**C**) Pathogen DNA accumulation in roots at 21 dpi. Individual points represent biological replicates. (**D-E**) Representative micrographs showing *P. brassicae* spores within root hairs of Col-0 and Col-0::35S*Pb*GH_3_ line 14.4 at 3 dpi (scale bars = 50 µm). (**F**) Number of infected root hairs at 3 dpi in Col-0 and Col-0::35S*Pb*GH_3_ lines (n = 20). For B, C, and E statistical significance relative to Col-0 is indicated (*p* < 0.05; ns, not significant).

Analysis of *Pb*GH_3_ transcript abundance in Col-0 from 0 to 21 dpi revealed that *Pb*GH_3_ is expressed during primary infection, peaking at 2 dpi (**Figure S8, Table S10**). Consistent with the increased root hair density observed in *Pb*GH_3_ overexpressing lines, transgenic plants exhibited enhanced root hair colonization by *P. brassicae* at 3 dpi (**Figure 4D–E, Figure S9, Table S11**). The results suggest that the early *Pb*GH_3_ expression could be promoting pathogen entry and initial colonization but does not determine the final disease outcome, which, based on literature, is likely influenced by additional host and pathogen factors acting at later stages of infection (Malinowski et al., 2019).

### *Pb*GH_3_ is sequence unrelated but structurally similar to *Arabidopsis* GH_3._12/PBS3

Following the phenotypic characterization and the absence of strong effects on auxin conjugation, we decide to further investigate the molecular function of *Pb*GH_3_. To this end, phylogenetic and structural analyses were performed to assess the evolutionary relationship of *Pb*GH_3_ with plant GH_3_ proteins and to identify possible functional similarities. Phylogenetic analysis revealed that *Pb*GH_3_ does not cluster with plant GH_3_ proteins, indicating that it is not closely related to Arabidopsis GH_3_ family members at the sequence level (**Figure 5A, Figure S10**). In contrast, structural comparisons between the predicted *Pb*GH_3_ protein and *Arabidopsis* GH_3_ proteins revealed a high degree of structural similarity with *At*GH_3.12_/PBS3) and *At*GH_3.7_, both members of clade III (**Figure 5B**). In addition, a third *Arabidopsis* GH_3_-like protein (*AT*5g13380) showed structural similarity with *Pb*GH_3_, although this protein has not yet been functionally characterized (**Figure 5B**). Given that clade III GH_3_ proteins are primarily associated with SA metabolism rather than auxin conjugation (Nobuta et al., 2007; Okrent et al., 2009), the observed structural similarity suggests that *Pb*GH_3_ may participate in SA-related metabolic processes.

**FIGURE 5:**
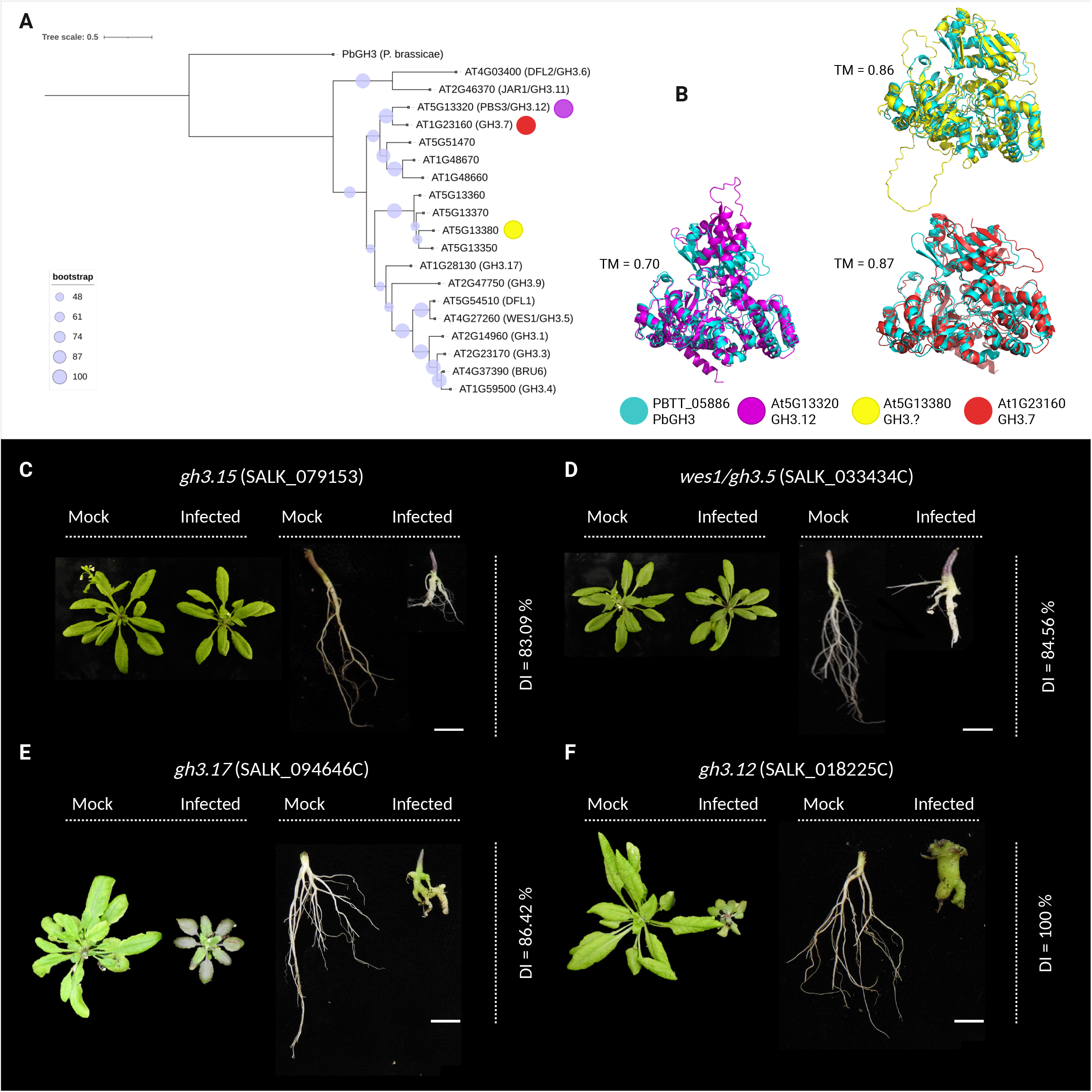
Structural similarity between *Pb*GH_3_ and *Arabidopsis At*GH_3.12_/PBS3 is linked to increased clubroot susceptibility in *gh3*.*12* mutants. (**A**) Phylogenetic analysis of *Pb*GH_3_ and *Arabidopsis* thaliana GH_3_ family members. The maximum-likelihood tree was inferred using IQ-TREE2 with the LG+I+G4 model (BIC-selected), with bootstrap support values (1000 ultrafast replicates) represented by circles at internal nodes proportional to support value. Branch lengths are drawn to scale (scale bar = 0.5 substitutions per site). (**B**) Structural superposition of *Pb*GH_3_ with selected *Arabidopsis* GH_3_ proteins. *Pb*GH_3_ (cyan) shows high structural similarity with *At*GH_3.12_/PBS3 (magenta) and *At*GH_3.7_ (red), with TM-scores indicated for each comparison. A third *Arabidopsis* GH_3.?_ protein (*At*5g13380, yellow) also shows partial structural alignment. (C–F) Representative images of GH_3_ mutants *Arabidopsis* plants inoculated with water or *P. brassicae* pathotype 3A at 21 dpi. Mutants include gh3.15 (**C**), wes1/gh3.5 (**D**), gh3.17 (**E**), and *gh3*.*12*/pbs3 (**F**). Whole rosettes and corresponding root systems are presented. DI values are indicated for each genotype. Scale bars = 0.1 cm.

### Loss of function of the gene GH_3.12_ /PBS3 increases clubroot susceptibility

Given the structural similarity between *Pb*GH_3_ and GH_3.12_/PBS3, we next investigated whether the disruption of GH_3.12_/PBS3 affects the progression of clubroot infection. In parallel, additional GH_3_ knockout mutants for other *Arabidopsis* GH_3_ belogning to clade I and II (*gh3*.*5, gh3*.*15*, and *gh3*.*17*) were included as controls to evaluate if altered clubroot susceptibility is specific to the loss of GH_3.12_/PBS3 or reflects a broader contribution of GH_3_ family members.

The mutant lines of *gh3*.*5, gh3*.*15*, and *gh3*.*17* displayed clubroot susceptibility levels similar to the WT Col-0, with no significant differences in disease index (DI) (**Figure 5C–E, Table S12**). In contrast, and consistent with its unique clade III identity and established role in SA metabolism (Nobuta et al., 2007), *gh3*.*12* mutants exhibited a pronounced increase in clubroot susceptibility, reaching a disease index of 100% (**Figure 5F**). Together, these results provide direct evidence for the importance of GH_3.12_/PBS3 in resistance to *P. brassicae* and further support an indirect functional link between *Pb*GH_3_ and SA metabolism.

### *Pb*GH_3_ affects SA-JA balance *in planta*

Given the role of SA in regulating root development and the structural similarity of *Pb*GH_3_ to GH_3.12_/PBS3, a key enzyme in SA metabolism, endogenous SA and JA levels were quantified in *Pb*GH_3_ overexpressing *Arabidopsis* lines and compared with Col-0. Hormone profiling revealed no significant differences in free and conjugated SA levels between Col-0 and the three transgenic lines in the case of SA; however, SA conjugates were significantly higher in line 14.4 (**Figure 6A–B, Table S13**). In contrast, jasmonate profiling showed a consistent reduction in JA levels in *Pb*GH_3_ overexpressing lines relative to Col-0 (**Figure 6C**). The bioactive conjugate JA–Ile was detected in Col-0 but was undetectable in *Pb*GH_3_ transgenic lines (**Figure 6C, Table S13**). Together, these results showed that the expression of *Pb*GH_3_ alters jasmonate homeostasis suggesting an indirect effect on SA metabolism through the antagonism between SA and JA. This modulation of SA- and JA-related hormones supports a role for *Pb*GH_3_ in modulating the host hormonal balance during infection.

**FIGURE 6:**
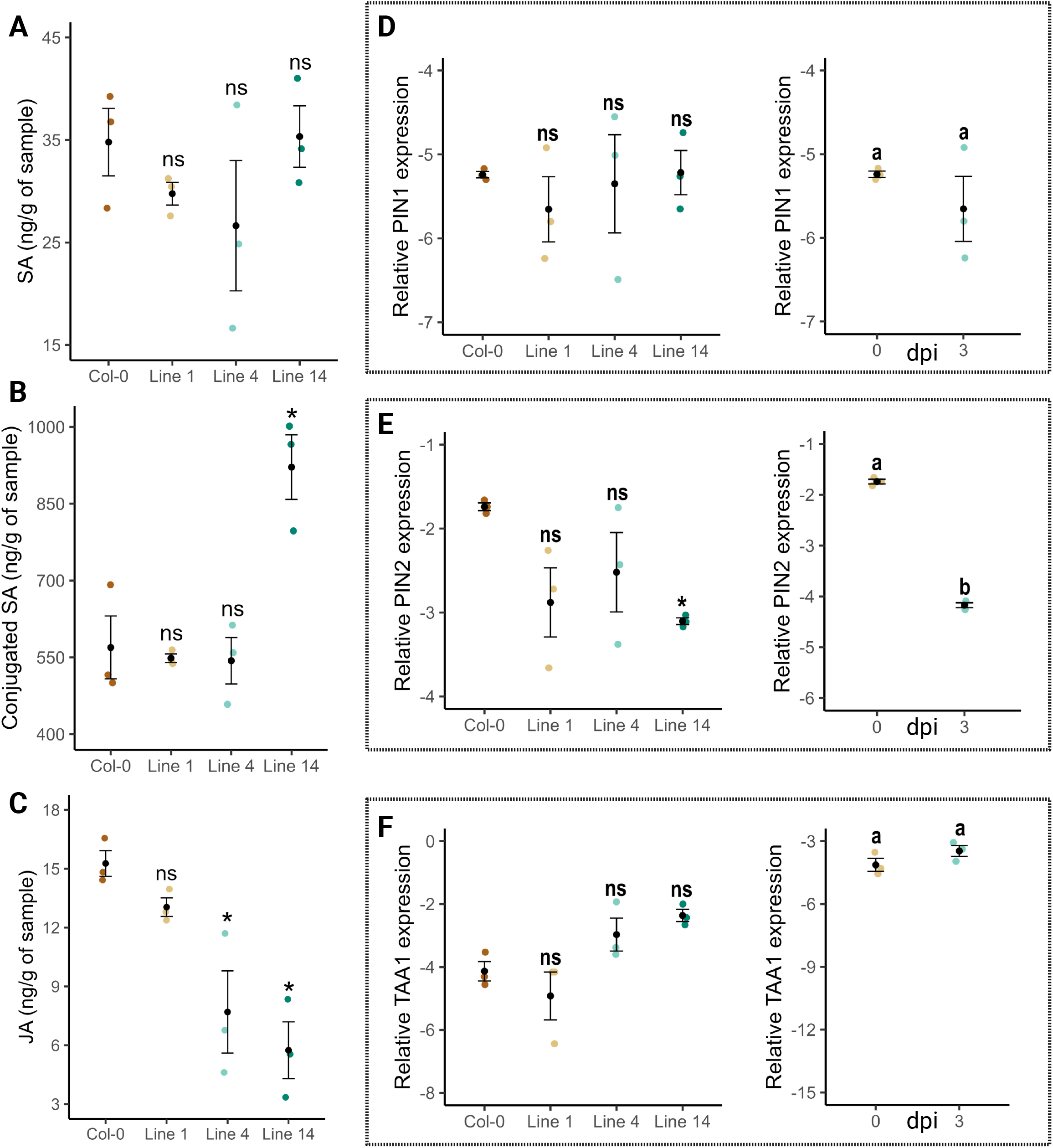
*Pb*GH_3_ overexpression alters SA–JA homeostasis and induces transcriptional responses associated with early clubroot infection. (**A**) Quantification of free salicylic acid (SA) levels in roots of wild-type Col-0 and Col-0::35S*Pb*GH_3_ *Arabidopsis* lines at 20 dpg. (**B**) Levels of conjugated SA in the same genotypes. (**C**) Jasmonic acid (JA) content in roots of Col-0 and Col-0::35S*Pb*GH_3_ lines at 20 dpg. (D–F) Relative expression levels of auxin-related marker genes PIN_1_ (**D**), PIN_2_ (**E**) and TAA_1_ (**F**) in Col-0 and Col-0::35S*Pb*GH_3_ lines under non-infected conditions (left panels) and during early Plasmodiophora brassicae infection at 3 dpi (right panels). Individual data points represent biological replicates. Asterisks indicate significant differences relative to Col-0 (*p* < 0.05); ns, not significant. Different letters designate statistically significant differences between time points (0 vs 3 dpi; *p* < 0.05).

### *Pb*GH_3_ induces expression responses consistent with early clubroot infection

Despite the lack of evidence for a direct role in auxin conjugation, our previous results showed that *Pb*GH_3_ overexpression induces root architectural changes that facilitate early clubroot infection. Given the central role of auxin as a master regulator of root development (Saini et al, 2013) these observations suggest that *Pb*GH_3_ may indirectly influence auxin-related regulatory pathways. To explore whether these alterations are reflected at the transcriptional level, key components of polar auxin transport and biosynthesis were examined. Expression of *PIN*_*1*_ was unchanged among genotypes and was not affected during early infection at 3 dpi (**Figure 6D, Table S14**). In contrast, *PIN*_*2*_ expression was slightly reduced in all *Pb*GH_3_ overexpressing lines, being line 14.4 the only one that showed a significant repression consistent with the downregulation detected in Col-0 during early clubroot disease at 3 dpi (**Figure 6E, Table S14**). The auxin biosynthetic marker *TAA*_*1*_ showed no significant differences in expression between the Col-0::35S*Pb*GH_3_ lines and Col-0, nor during primary infection (**Figure 6F, Table S14**). Together, these data suggest that *Pb*GH_3_ overexpression does not affect auxin biosynthesis but may alters PIN-mediated basipetal auxin transport, a process known to regulate auxin distribution and accumulation in roots (Blilou et al., 2005; Vanneste & Friml, 2009), and known to take place at early stages of clubroot development and gall initiation (Siemens et al., 2006; Ludwig-Müller et al., 2011).

### *Pb*GH_3_ partially complement *At*GH_3.12_/PBS3 lost in *A. thaliana*

Based on the structural similarity between *Pb*GH_3_ and GH_3.12_/PBS3, we cannot discard that *Pb*GH_3_ had a function analogous to that of GH_3.12_/PBS3 in SA biosynthesis. To test this, *gh3*.*12* mutant plants and *gh3*.*12*::35S*Pb*GH_3_ lines were analyzed to determine whether *Pb*GH_3_ expression can complement the loss of function of GH_3.12_/PBS3 in root and leaf development and clubroot disease progression. Integration and expression of the *Pb*GH_3_ transgene in the *gh3*.*12* background were confirmed by PCR and RT–qPCR, respectively (**Table S15**).

We observed that *gh3*.*12*::35S*Pb*GH_3_ plants exhibited epinastic leaves similar to Col-0::35S*Pb*GH_3_ lines, whereas the *gh3*.*12* was phenotypically identical to the wild type Col-0 (**Figure 7A, C**). The analysis of the root morphology showed that both *gh3*.*12* and *gh3*.*12*::35S*Pb*GH_3_ lines had a significantly higher number of root hairs compared to Col-0 and Col-0::35S*Pb*GH_3_ (**Figure 7A–B, Table S16**). However, this increased in root hairs was lower when compared *gh3*.*12*::35S*Pb*GH_3_ lines to *gh3*.*12* mutant (**Figure 7A–B**). Other developmental traits, including hypocotyl length, primary root length, and lateral root number, did not differ significantly among genotypes (**Table S16**). To determine whether *Pb*GH_3_ overexpression affects clubroot susceptibility in the *gh3*.*12* background, *gh3*.*12*::35S*Pb*GH_3_ lines were inoculated with *P. brassicae* pathotype 3A and disease development was assessed at 21 dpi. All three independent *gh3*.*12*::35S*Pb*GH_3_ lines exhibited lower susceptibility, (DI= 85–95%) compared with the *gh3*.*12* mutant (DI = 100%). Relative to Col-0, only one *gh3*.*12*::35S*Pb*GH_3_ line displayed increased susceptibility (**Figure 7C–D, Table S17**).

**FIGURE 7:**
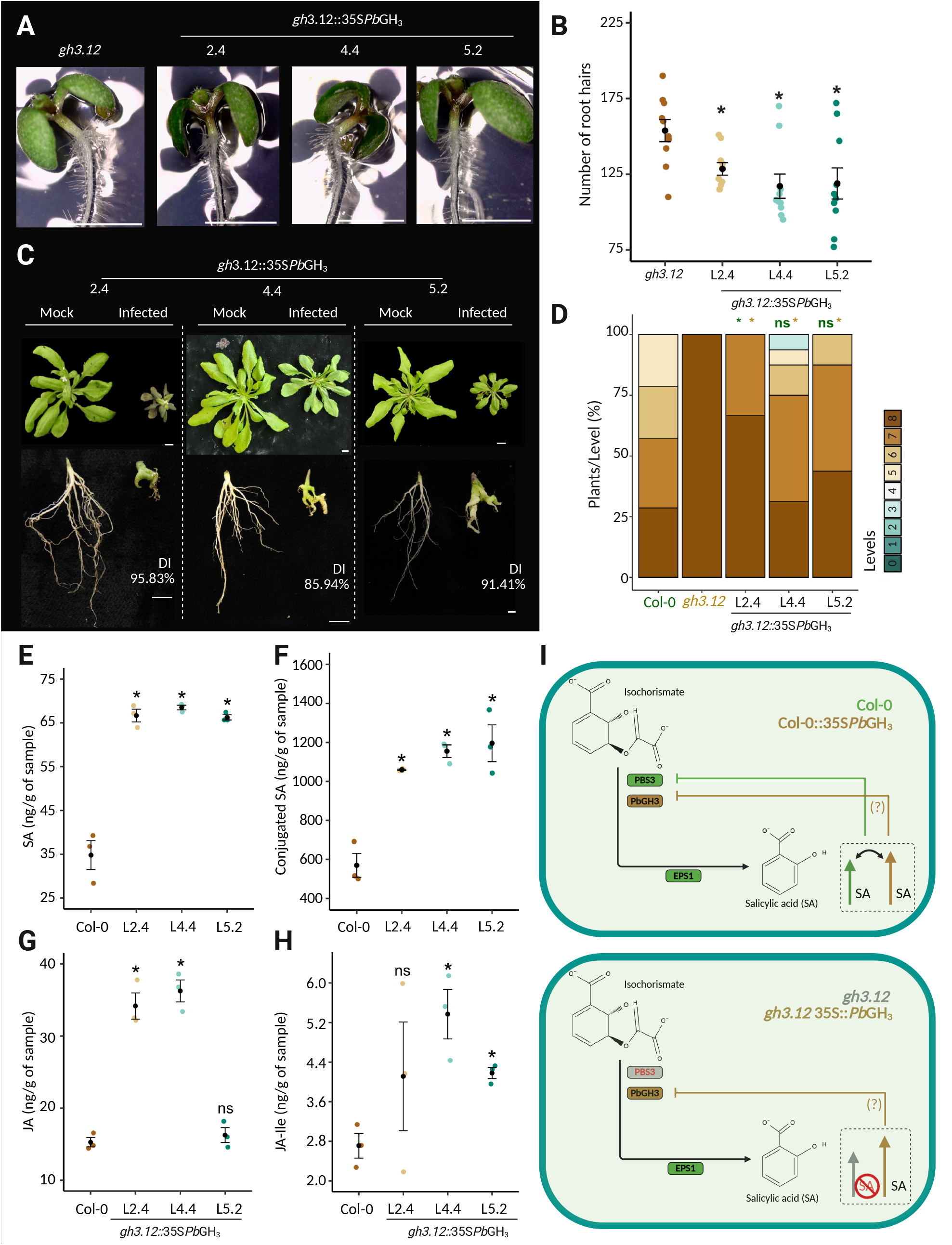
*Pb*GH_3_ partially complements the loss of *At*GH_3.12_/PBS3 by modulating root hair development, hormone homeostasis and clubroot susceptibility. (**A**) Representative images of primary roots from the *gh3*.*12* mutant and independent *gh3*.*12*::35S*Pb*GH_3_ lines 2.4, 4.4 and 5.2 at 10 dpg, illustrating reduced root hair density in complemented lines relative to *gh3*.*12* (scale bars = 1 cm). (**B**) Quantification of root hair number per primary root. Individual points represent single seedlings. (**C**) Representative images of *gh3*.*12*::35S*Pb*GH_3_ lines inoculated with water or *P. brassicae* pathotype 3A at 21 dpi. Whole rosettes and corresponding root systems are shown. DI values are indicated for each line (scale bars = 1 cm). (**D**) Distribution of disease severity levels at 21 dpi expressed as the percentage of plants per category. (**E**) Quantification of SA, conjugated SA (**F**), jasmonic acid (JA) (**G**) and JA–Ile (**H**) levels in roots of Col-0 and *gh3*.*1::*35S*Pb*GH_3_ lines at 20 dpg. (**I**) Working model illustrating the proposed role of *Pb*GH_3_ in the PBS3-dependent salicylic acid pathway. *Pb*GH_3_ partially compensates for the loss of *At*GH_3.12_/PBS3, contributing to the modulation of SA homeostasis and downstream hormonal balance, resulting in intermediate phenotypes relative to Col-0 (*p* < 0.05); ns, not significant.*gh3*.*12* and wild-type Col-0 plants. For panels **B** and **E**–**H**, asterisks indicate significant differences

Given the well-established reduction of SA accumulation in *gh3*.*12* mutants resulting from the loss of *PBS3/AtGH*_*3*.*12*_ function (Nobuta et al., 2007; Okrent et al., 2009), our hormone measurements focused on Col-0 and *gh3*.*12*::35S*Pb*GH_3_ lines. Hormone profiling revealed that *gh3*.*12*::35S*Pb*GH_3_ lines accumulated higher levels of SA and SA conjugates compared to Col-0, as well as increased levels of JA and JA–Ile (**Figure 7E–H, Table S18**). Coordinated activation of the SA and JA pathways has been reported during ETI through a non-canonical mechanism (Liu et al., 2016; Betsuyaku et al., 2018), suggesting that *Pb*GH_3_, in the absence of GH_3.12_/PBS3, may trigger these defense pathways. But this hypothesis requires further investigation. Although overexpression of *PbGH*_*3*_ in the *gh3*.*12* background caused a reduction in clubroot susceptibility and root hair abundance, *gh3*.*12*::35S*Pb*GH_3_ lines consistently displayed intermediate phenotypes between *gh3*.*12* and Col-0, both at the morphological and hormonal levels. These results suggest that *Pb*GH_3_ can partially restore hormone homeostasis and disease resistance in the absence of *At*GH_3.12_/PBS3 but not fully.

Together, we propose a model in which *Pb*GH_3_ may contribute to a tight regulation of SA homeostasis, potentially through feedback control of SA biosynthesis or preventing increases in free SA in the *Pb*GH_3_ overexpressing lines (**Figure 7I**). We then hypothesize that loss of GH_3.12_/PBS3 results in a collapse of SA accumulation and defense competence, whereas expression of *Pb*GH_3_ provides partial functional compensation, restoring hormone levels but without fully re-establishing wild-type regulation. This altered SA homeostasis may be linked to the auxin-associated developmental phenotypes observed in the 35S*Pb*GH_3_ lines, including changes in root architecture (Wang et al., 2007; Pasternak et al., 2019).

## DISCUSSION

Previous studies have established that *P. brassicae* actively manipulates host hormone metabolism to promote infection, particularly through auxin and SA pathways. While the SABATH-type methyltransferase *Pb*BSMT suppresses local immunity by converting SA into inactive methyl salicylate (MeSA) (Ludwig-Müller et al., 2015; Benade et al., 2025), other pathogens are known to interfere directly with SA biosynthesis (Djamei et al., 2011; Bauters et al., 2020). Although *Pb*GH_3_ was proposed as a candidate effector involved in auxin conjugation (Schwelm et al., 2015) and has been shown *in vitro* to conjugate auxin to multiple amino acids (Schwelm et al., 2015; Smolko et al., 2024), its *in planta* activity was only recently investigated, and the effector’s role remains inconclusive (Smolko et al., 2024). Our results provide functional evidence that *Pb*GH_3_ contributes to disease-associated phenotypes by altering host hormone homeostasis, supporting its role as a virulence factor.

Constitutive expression of *Pb*GH_3_ in *Arabidopsis* resulted in developmental alterations, including reduced apical dominance, epinastic leaves, enhanced lateral branching, and increased root hair formation. These *Pb*GH_3_-associated phenotypes were conserved in canola, where transgenic lines displayed reduced apical dominance and a significantly higher number of root hairs relative to the WT cultivar, indicating that *Pb*GH_3_ affects regulatory pathways conserved across Brassicaceae. Such traits are classically associated with perturbed auxin distribution rather than global changes in auxin abundance (Grieneisen et al., 2007; Bennet et al., 2010). Apical dominance is controlled by polar auxin transport from the shoot apex via PIN-mediated efflux (Leyser, 2005; Balla et al, 2016), epinasty reflects asymmetric auxin accumulation in leaf tissues (Sandalio et al., 2016), and root hair initiation and elongation depend on tightly regulated local auxin maxima in epidermal cells (Jones et al., 2009). Together, these phenotypes alterations are consistent with a model in which *Pb*GH_3_ disrupts auxin gradients between shoot and root, leading to relative auxin accumulation in roots and depletion in aerial tissues, although these changes were not significative (**Figure 2**).

Multiple evidences argue against a direct role for *Pb*GH_3_ in auxin conjugation *in planta* (**Figure 2**; Smolko et al., 2024). Auxin levels and conjugate profiles did not change between Col-0 and Col-0::35S*Pb*GH_3_, *Pb*GH_3_ overexpressing plants did not exhibit increased tolerance to exogenous auxin, and responses to synthetic auxins were not substantially altered. These observations exclude *Pb*GH_3_ as a functional auxin-amido synthetase *in vivo* and instead indicate that the auxin-associated developmental phenotypes are an indirect consequence of *Pb*GH_3_ expression. A more suitable explanation is perturbation of SA-dependent regulation of auxin transport and signaling, as SA is known to modulate polar auxin transport through effects on PIN abundance, localization, and activity (Pasternak et al., 2019; Tan et al., 2019; Zhou et al., 2022). Such SA-mediated modulation of auxin gradients provides a mechanistic framework linking altered hormone homeostasis to the observed changes in shoot architecture and root hair development.

*Pb*GH_3_ expression peaks early during infection, coinciding with the primary infection phase characterized by root hair penetration, a pattern consistent with previous RNA-seq data (Schwelm et al., 2015). A more recent study using semi-quantitative PCR, however, reported high expression at 28 dpi, so late in the disease cycle that could reflect re-infection events initiated by newly released single spores (Liu et al., 2020). This timing and strict transcriptomic regulation are consistent with a role in facilitating pathogen entry rather than determining later stages of gall development, which depend on additional host and pathogen factors (Kageyama & Asano, 2009; Hwang et al., 2015; Liu et al., 2020, Pérez-López et al., 2019; 2020; Mukhopadhyay et al., 2025). Increased root hair formation has been documented across diverse plant–microbe interactions and is frequently associated with hormone crosstalk involving SA, JA, ethylene, and auxin transport (Vissenberg et al., 2020; Pecenková et al., 2017; Jan et al., 2024). Importantly, enhanced root hair infection does not necessarily translate into increased disease severity, as primary infection alone is insufficient for successful clubroot development (Yang et al., 2022). The limited impact of *Pb*GH_3_ overexpression on final disease outcome therefore aligns with current models of clubroot progression in which early colonization and gall development are mechanistically decoupled (Kageyama & Asano, 2009; Liu et al., 2020; Muirhead and Pérez-López, 2022; Javed et al., 2023).

Structural similarity between *Pb*GH_3_ and Brassicaceae GH_3_ group III proteins, particularly GH_3.12_/PBS3 and GH_3.7_, places *Pb*GH_3_ within a functional role associated with SA metabolism rather than auxin conjugation. GH_3.12_/PBS3 plays a central and non-redundant role in SA biosynthesis in *Arabidopsis* by catalyzing the formation of isochorismate–glutamate, a key precursor in pathogen-induced SA accumulation (Nobuta et al., 2007; Okrent et al., 2009). Recent comparative analyses indicate that this PBS3-dependent SA pathway is largely restricted to Brassicales and emerged through lineage-specific duplication and functional specialization of GH_3_ genes, together with coordinated evolution of transport components (Hong et al., 2025).

The partial complementation of GH_3.12_/PBS3 loss by *Pb*GH_3_ provides important insight into the mode of action of this effector. Expression of *Pb*GH_3_ in the *gh3*.*12* mutant background restored SA-related hormonal responses and reduced disease susceptibility relative to the mutant, yet developmental and immune phenotypes were not fully reverted to WT levels. This is consistent with functional overlap within the same regulatory pathway rather than full enzymatic equivalence. GH_3.12_/PBS3 operates within a highly regulated SA biosynthetic module, and small changes in its activity or regulation have pronounced effects on immune competence and development (Nobuta et al., 2007; Okrent et al., 2009). Such partial complementation is a recurrent feature of pathogen effectors that act as structural or functional mimics of host proteins. Rather than reproducing full host activity, these effectors often restore only a subset of biochemical or regulatory functions, allowing pathogens to perturb host signaling without triggering excessive defense activation (**Figure 7I**). Well-characterized examples include phytoplasma effectors that structurally mimic transcriptional regulators yet lack full DNA-binding or regulatory capacity, resulting in altered developmental programs rather than complete pathway takeover (MacLean et al., 2011; MacLean et al., 2014; Yang et al., 2025). Similarly, bacterial and oomycete effectors that resemble host kinases or ubiquitin ligases frequently act as modulators rather than functional replacements, reshaping signaling outputs (Mukhtar et al., 2011; Djamei et al., 2011). *Pb*GH_3_ fits within this paradigm by acting as a partial functional mimic of GH_3.12_/PBS3 that perturbs SA balance without fully reproducing GH_3.12_/PBS3’s biological function.

The requirement for such strict regulation of *Pb*GH_3_ during the disease is particularly evident given the central role of SA in plant immunity. Sustained activation of SA biosynthesis is associated with developmental penalties and reduced host fitness and is therefore unlikely to be advantageous for obligate biotrophic pathogens that require prolonged host viability (Huot et al., 2014; Karasov et al., 2017; Chen et al., 2017; Mine et al., 2018). The observation that *Pb*GH_3_ expression does not induce SA accumulation, even when overexpressed, suggests that its activity is limited or tightly regulated. *Pb*GH_3_ may contribute to a buffering or competitive interaction with GH_3.12_/PBS3, altering SA conjugation and downstream signaling without triggering a full immune response that would compromise infection. An additional layer of control may reside in the delivery and localization of *Pb*GH_3_. Unlike many characterized plant pathogen effectors, *Pb*GH_3_ lacks a canonical N-terminal signal peptide (**Figure S11**), raising the possibility that it is secreted via a non-classical pathway yet to be discovered or delivered at low efficiency into host cells. Non-canonical secretion has been reported for several microbial effectors and is increasingly recognized as a strategy to limit effector abundance or spatial distribution within host tissues (Liu et al., 2014; Wang et al., 2017). For *Pb*GH_3_, such a mechanism could further restrict its effective concentration in host cells, preventing excessive perturbation of SA metabolism while still allowing localized or transient interference during early infection stages.

## CONCLUSIONS

This study provides evidence that the *P. brassicae* effector *Pb*GH_3_ is not a canonical auxin- or JA-conjugating enzyme when expressed *in planta*, however it induces conserved auxin-like developmental phenotypes and root changes in *Arabidopsis* and canola. *Pb*GH_3_ overexpression *in planta* promotes root hair formation favoring early root hair colonization by *P. brassicae*. Genetic and structural evidence links *Pb*GH_3_ to *Arabidopsis* clade III GH_3.12_/PBS3 function mediating SA biosynthesis pathway. The partial complementation induced by overexpression of *Pb*GH_3_ in *gh3*.*12* background supports a functional interaction with this pathway rather than direct replacement of GH_3.12_/PBS3 activity.

Together, our results support a model in which *Pb*GH_3_ modulates SA homeostasis and downstream hormonal crosstalk, associated with *PIN*_*2*_ repression which can supports an altered auxin redistribution, to create a root environment permissive for early colonization. Because *Pb*GH_3_ overexpression alone does not consistently increase final disease severity, these findings further highlight that clubroot progression depends on additional pathogen factors and later-stage host– pathogen interactions. *Pb*GH_3_ therefore represents a key component of hormonal manipulation in clubroot pathogenesis and a potential target for durable disease control strategies.

## ACKNOWLEDGEMENTS

This work was funded by the Canola Agronomic Research Program (Grant ID 2021.4), Western Grain Research Foundation, Canola Council of Canada, Alberta Canola, and Manitoba Canola Growers Association, as well as by the Discovery Program (Grant ID RGPIN-2021–02518) of the Natural Sciences and Engineering Research Council of Canada. We are also thankful to the Fonds de Recherche du Québec – Nature et technologies division, and the Faculté des Sciences de l’Agriculture et de l’Alimentation at Université Laval, for supporting MGG through a doctoral scholarship.

## AUTHOR CONTRIBUTIONS

EPL conceived and funded the project. EPL, MS, EF, and IM supervised and guided the execution of the experiments. MGG, MSV, JW, SM, EF, RM, KS, and EPL performed experiments and analyzed the data. IM provided the infrastructure required for phenotyping. MSV and JW generated the Arabidopsis and canola overexpression lines, respectively. RM and KS conducted vascular morphology analyses. SM performed AlphaFold structural analyses. MGG, KS, and EPL prepared the figures and tables. MGG and EPL wrote the manuscript with contributions from all authors.

## DATA AVAILABILITY

The data that support the findings of this study are available within the article and in the Supporting Information (Figs S1–S10 and Tables S1–S18). The plasmid constructed in this study and used to transform *A. thaliana* and *B. napus* was deposited to Addgene (plasmid #232335; RRID: Addgene_232335).

## SUPPORTING INFORM*AT*ION SUPPLEMENTARY TABLES

**TABLE S1**. Primers used for qPCR, RT–qPCR and PCR analyses.

**TABLE S2**. *Pb*GH_3_ expression levels in *Arabidopsis thaliana* transgenic lines in the Col-0 background.

**TABLE S3**. *Pb*GH_3_ expression levels in *Brassica napus* Westar cv. transgenic lines.

**TABLE S4**. Above-ground phenotypic traits of *Pb*GH_3_ overexpressing *Arabidopsis thaliana* and *Brassica napus* lines.

**TABLE S5**. Primary root length of *Arabidopsis* seedlings in response to auxin treatments.

**TABLE S6**. Hormone profiling of Col-0::35S*Pb*GH_3_ *Arabidopsis thaliana* lines.

**TABLE S7**. Root phenotypic traits of *Pb*GH_3_ overexpressing lines.

**TABLE S8**. Individual clubroot disease severity scores used for disease index calculations in wild-type Col-0 and Col-0::35S*Pb*GH_3_ lines inoculated with *Plasmodiophora brassicae* pathotype 3A.

**TABLE S9**. Quantitative PCR cycle threshold (Ct) values for *Plasmodiophora brassicae* (*Pb18S*) and *Arabidopsis* reference gene *AtSK11* during clubroot infection.

**TABLE S10**. Quantitative PCR cycle threshold (Ct) values of the *Plasmodiophora brassicae* gene *Pb*GH_3_ during infection of *Arabidopsis* roots.

**TABLE S11**. Number of infected root hairs quantified at 3 days post-inoculation (dpi) in *Arabidopsis thaliana* Col-0 and Col-0::35S*Pb*GH_3_ transgenic lines 1.5, 4.1 and 14.4, following inoculation with *Plasmodiophora brassicae* pathotype 3A.

**TABLE S12**. Individual clubroot disease severity scores used for disease index calculations in GH_3_ *Arabidopsis* knock-out mutants inoculated with *Plasmodiophora brassicae* pathotype 3A.

**TABLE S13**. Salicylic acid and jasmonate levels in roots of wild-type Col-0 and Col-0::35S*Pb*GH_3_ Arabidopsis lines at 20 days post-germination (dpg).

**TABLE S14**. qRT–PCR Ct values for auxin-related genes in *Arabidopsis* Col-0 and Col-0::35S*Pb*GH_3_ lines.

**TABLE S15**. qRT–PCR Ct values of *Pb*GH_3_ expression in the *gh3*.*12*::35S*Pb*GH_3_ transgenic lines 2.2, 4.1 and 5.1.

**TABLE S16**. Root phenotypic measurements of *gh3*.*12* and *gh3*.*12*::35S*Pb*GH_3_ *Arabidopsis* lines 2.2, 4.1 and 5.1.

**TABLE S17**. Clubroot disease severity scores in *gh3*.*12*::35S*Pb*GH_3_ *Arabidopsis* lines and disease index (DI) following inoculation with *Plasmodiophora brassicae* pathotype 3A.

**TABLE S18**. Salicylic acid (SA), conjugated SA, jasmonic acid (JA) and JA-Ile conjugate levels in gh3.12 and *gh3*.*12*::35S*Pb*GH_3_ roots of *Arabidopsis* lines 2.2, 4.2, and 5.1 were evaluated at 20 days post-germination.

## SUPPLEMENTARY FIGURES

**FIGURE S1**. Addgene Full Sequence Map for pMDC32_ *Pb*GH_3_.

**FIGURE S2**. Validation of *Pb*GH_3_ transgene integration and expression in Col-0::35S*Pb*GH_3_ *Arabidopsis* and canola lines.

**FIGURE S3**. *Pb*GH_3_ overexpression does not affect petiole length or floral morphology but reduces silique length in *Arabidopsis*.

**FIGURE S4**. *Pb*GH_3_ overexpression does not alter auxin sensitivity during early seedling development.

**FIGURE S5**. *Pb*GH_3_ overexpression does not significantly affect cytokinin, ABA or PA levels in *Arabidopsis*.

**FIGURE S6**. *Pb*GH_3_ overexpression alters root architecture in *Brassica napus*.

**FIGURE S7**. Phenotype of *Arabidopsis thaliana* Col-0 at 21 days post-inoculation with *P. brassicae*.

**FIGURE S8**. Representative micrographs showing *P. brassicae* spores within root hairs of Col-0 and Col-0::35S*Pb*GH_3_ lines

**FIGURE S9**. Representative micrographs showing *P. brassicae* spores within root hairs of Col-0 and Col-0::35S*Pb*GH_3_ lines.

**FIGURE S10**. Sequence and structural conservation of *Pb*GH_3_ relative to *Arabidopsis* GH_3_ family members.

**FIGURE S11**. Predicted disordered N-terminal region of *Pb*GH_3_

