## Supplementary Figures for "A clubroot pathogen PBS3-like effector manipulates hormonal crosstalk to alter root morphology during colonization"

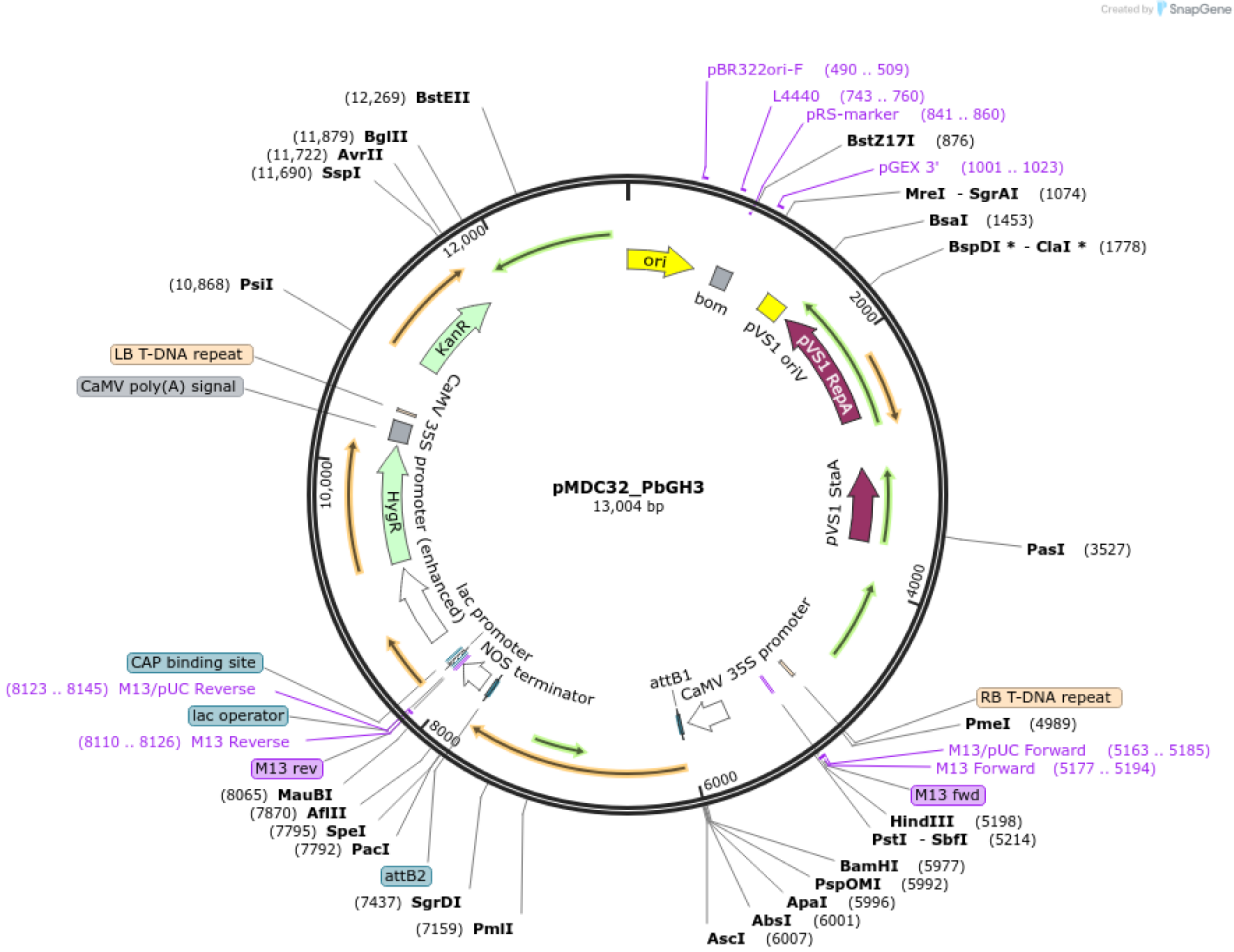


**FIGURE S1.** Addgene Full Sequence Map for pMDC32_ *Pb*GH$_3$.

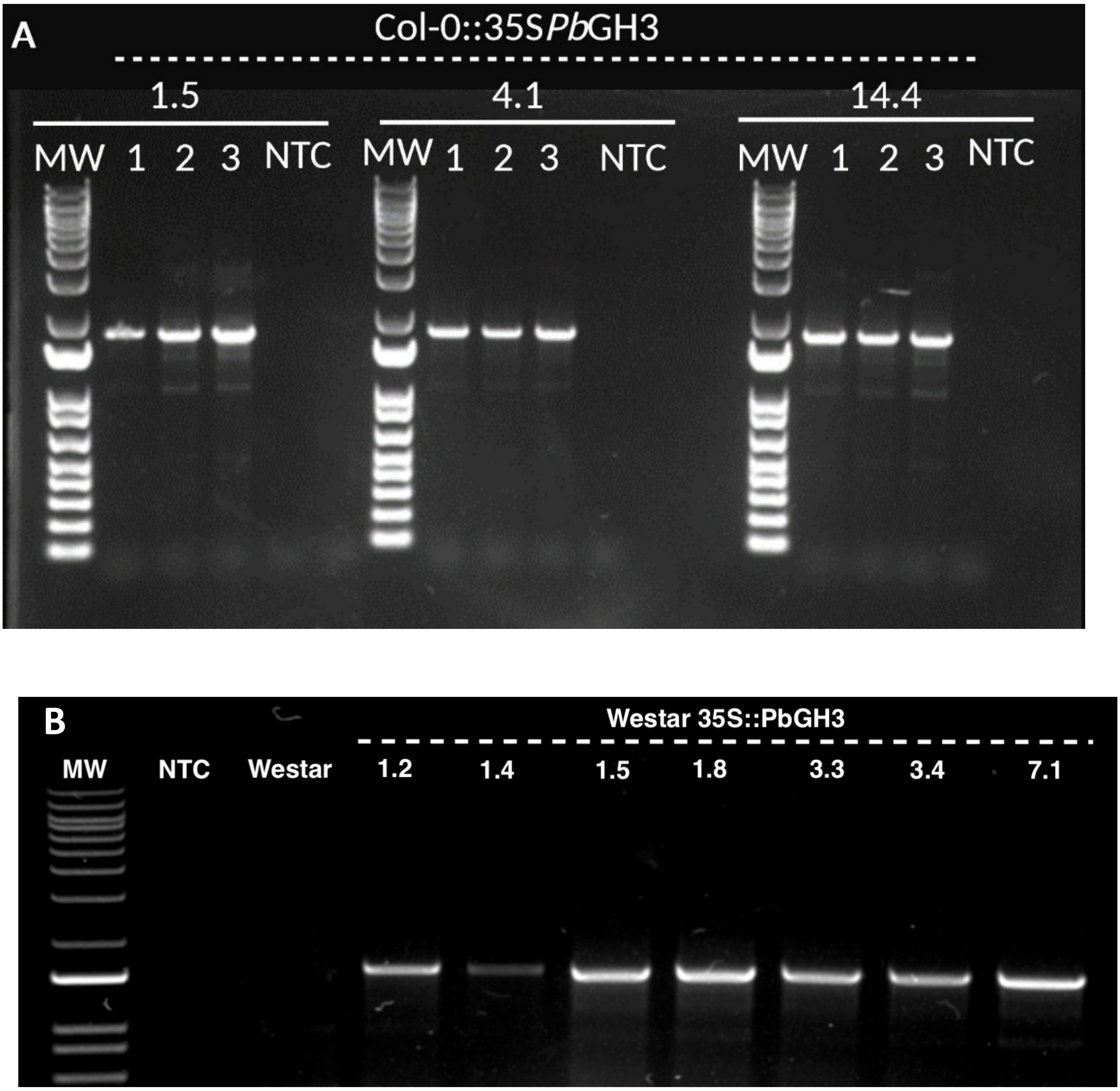


**FIGURE S2.** Validation of *Pb*GH$_3$ transgene integration in Col-0::35S *Pb*GH$_3$ *Arabidopsis* lines (A), and in canola lines (B).

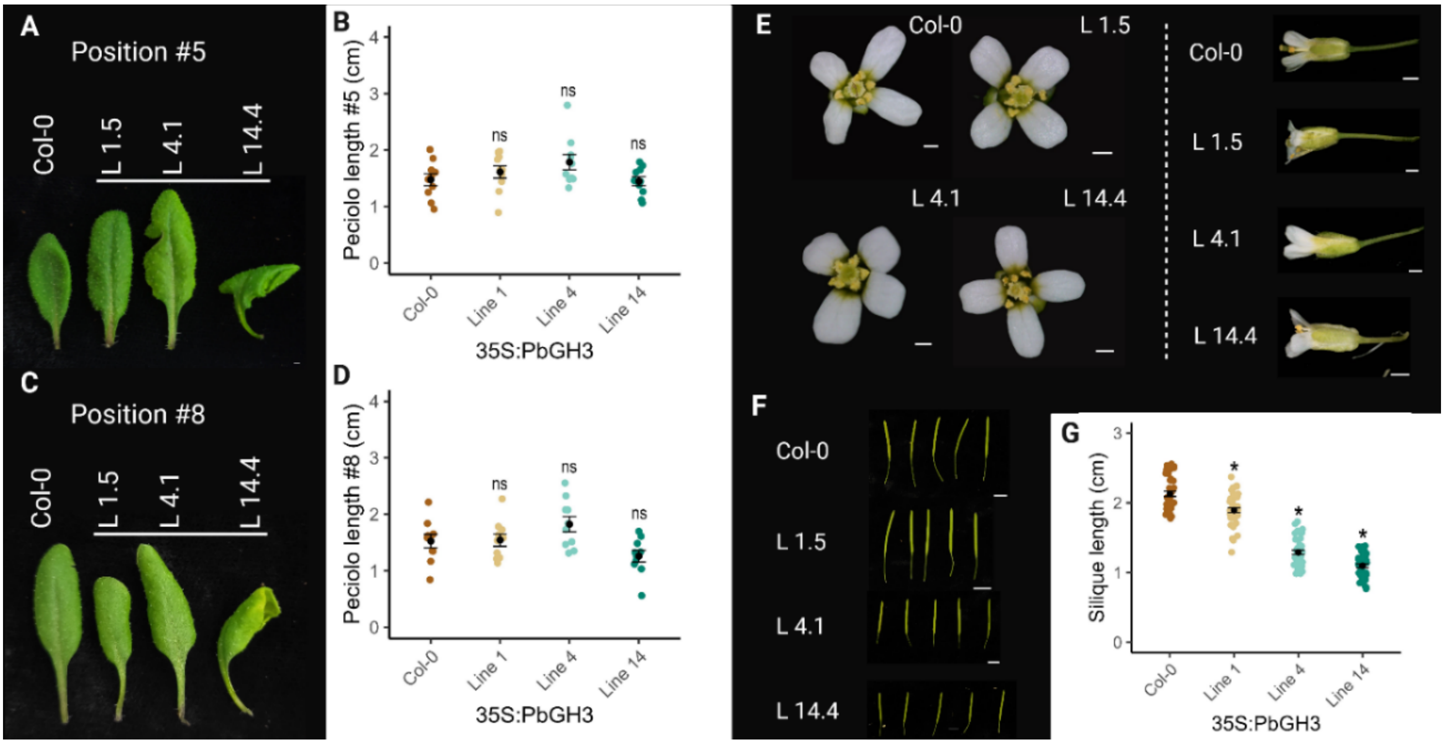


**FIGURE S3.** *Pb*GH3 overexpression does not affect petiole length or floral morphology but reduces silique length in *Arabidopsis*. (A, C) Representative images of rosette leaves at positions #5 (A) and #8 (C) from wild-type Col-0 and independent Col 35S::*Pb*GH3 *Arabidopsis* lines (L1.5, L4.1 and L14.4). (B, D) Quantification of petiole length at positions #5 (B) and #8 (D). No significant differences were detected between Col-0 and Col-0 35S::*Pb*GH3 lines. (E) Representative images of flowers and corresponding pistils from Col-0 and Col-0 35S::*Pb*GH3 lines, showing no overt floral defects. (F) Representative siliques from Col-0 and Col-0 35S::*Pb*GH3 lines. (G) Quantification of silique length, showing a significant reduction in Col-0::35S*Pb*GH3 lines compared with Col-0.

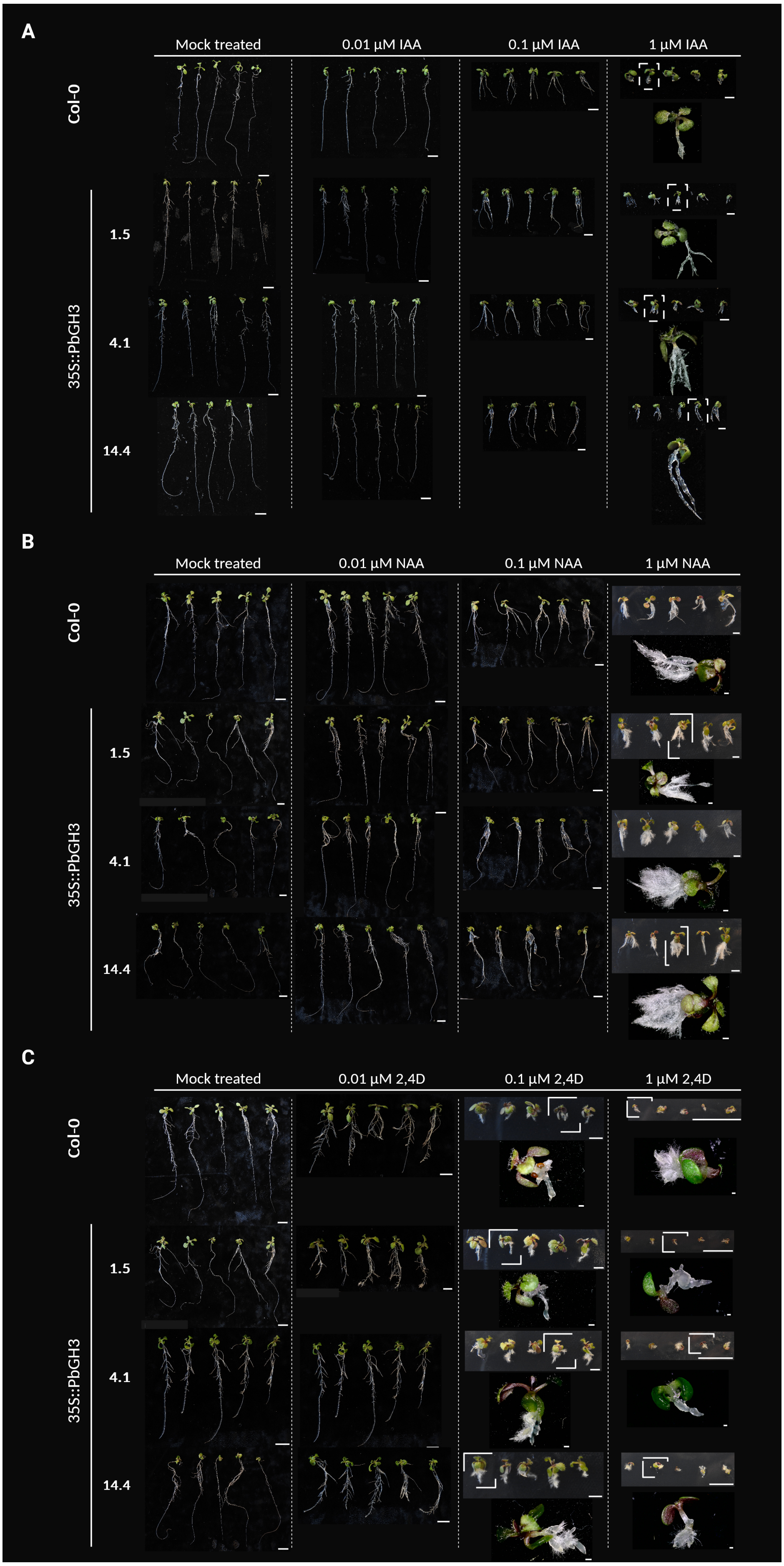

A
Mock treated
0.01 µM IAA
0.1 µM IAA
1 µM IAA
Col-0
35S::PbGH3
1.5
4.1
14.4
B
Mock treated
0.01 µM NAA
0.1 µM NAA
1 µM NAA
Col-0
35S::PbGH3
1.5
4.1
14.4
C
Mock treated
0.01 µM 2,4D
0.1 µM 2,4D
1 µM 2,4D
Col-0
35S::PbGH3
1.5
4.1
14.4

**FIGURE S4.** *Pb*GH$_3$ overexpression does not alter auxin sensitivity during early seedling development. Representative images of wild-type Col-0 and independent Col-0 35S::*Pb*GH$_3$ *Arabidopsis* lines (L1.5, L4.1 and L14.4) grown vertically on MS medium supplemented with increasing concentrations of auxins. (A) Seedlings treated with indole-3-acetic acid (IAA). (B) Seedlings treated with 2,4-dichlorophenoxyacetic acid (2,4-D). (C) Seedlings treated with naphthaleneacetic acid (NAA). Seedlings were imaged at 10 days post-germination. Comparable inhibition of primary root growth and overall seedling morphology were observed between Col-0 and Col-0 35S::*Pb*GH$_3$ lines across all auxin treatments. Scale bars = 1 cm.

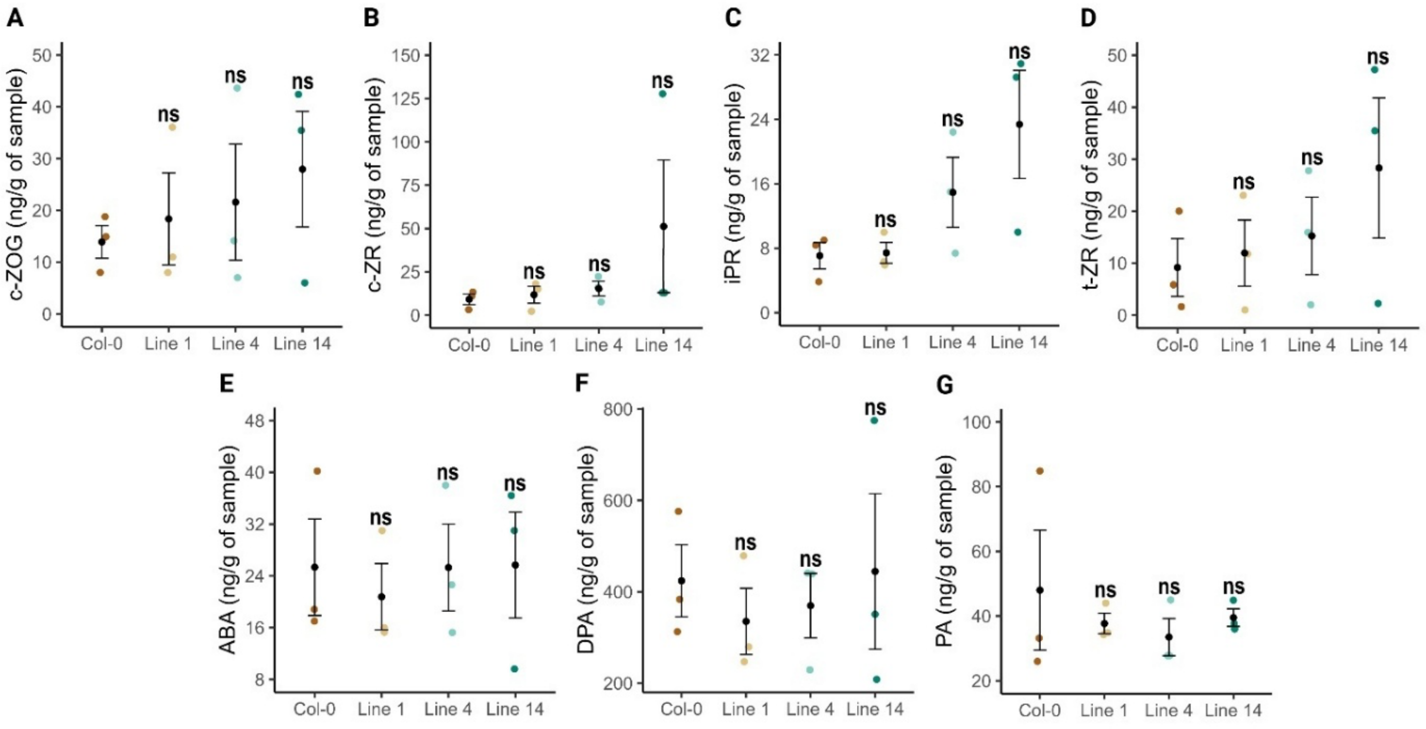


**FIGURE S5. *Pb*GH$_3$ overexpression does not significantly affect cytokinin, ABA or PA levels in *Arabidopsis*.** Quantification of endogenous hormone levels in wild-type Col-0 and independent Col-0 35S::*Pb*GH$_3$ *Arabidopsis* lines (Line 1, Line 4 and Line 14). (A) cis-zeatin-O-glucoside (cZOG), (B) cis-zeatin riboside (cZR),(C) isopentenyl riboside (iPR),(D) trans-zeatin riboside (tZR), (E) abscisic acid (ABA), (F) dihydrophaseic acid (DPA), and (G) phaseic acid (PA).

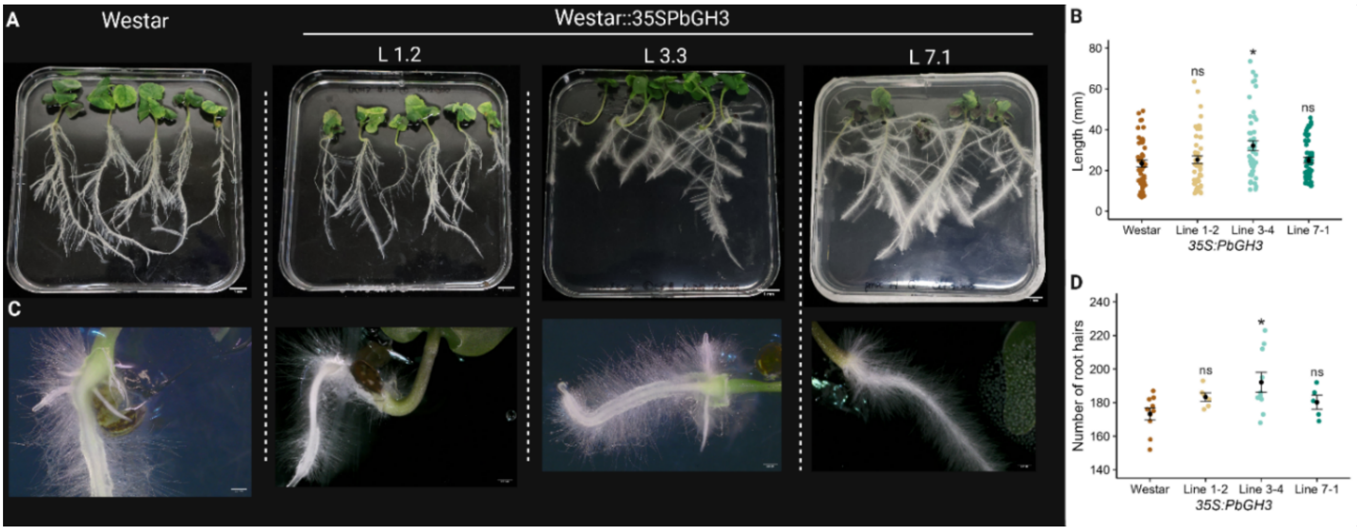


**FIGURE S6.** *Pb*GH3 overexpression alters root architecture in *Brassica napus*. (A) Representative images of root systems from wild-type Westar and independent Westar::35S*Pb*GH3 transgenic lines (L1.2, L3.3 and L7.1) grown under controlled conditions. (B) Quantification of primary root length in Westar and Westar 35S::*Pb*GH3 lines at 10 dpg . (C) Close-up views of root hair development in Westar and Westar 35S::*Pb*GH3 seedlings. (D) Quantification of root hair number per root at 5 dpg.

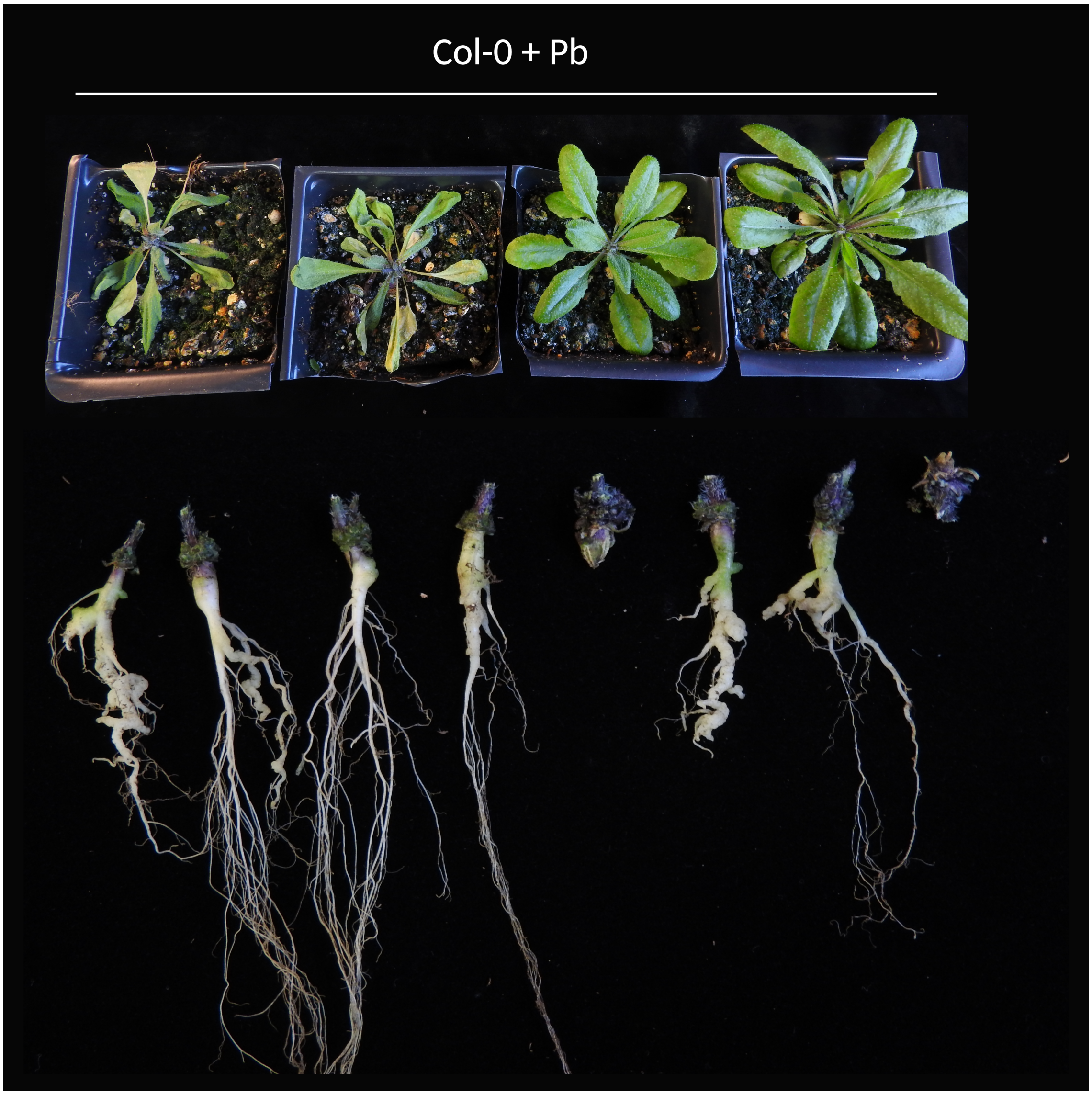


**FIGURE S7.** Phenotype of *Arabidopsis thaliana* Col-0 at 21 days post-inoculation with *P. brassicae*. The image illustrates the range of shoot phenotypes observed in inoculated plants.

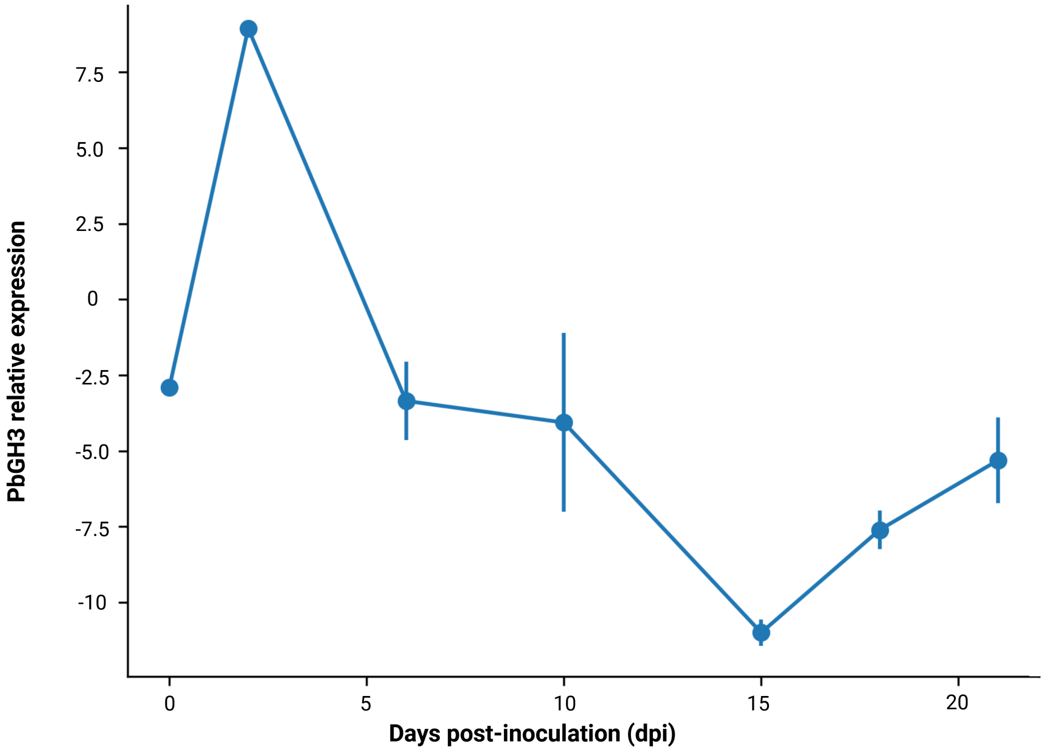


**FIGURE S8.** Expression profile of *PbGH3* during clubroot infection at 0, 2, 6, 10, 15, 18, and 21 dpi. Values represent the mean of three biological replicates, and bars indicate the standard error. The corresponding data are provided in Table S10.

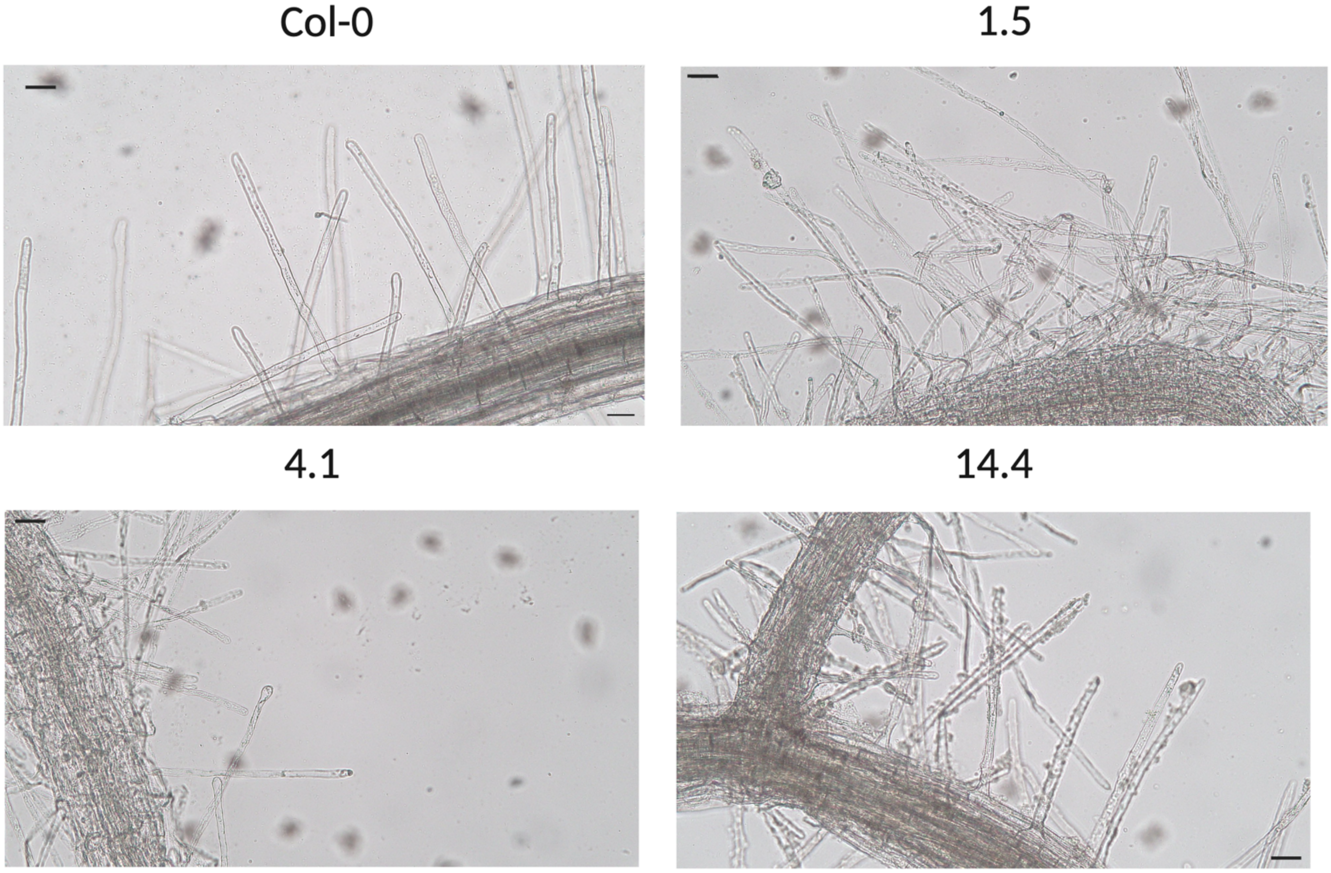


**FIGURE S9.** Representative micrographs showing *P. brassicae* spores within root hairs of Col-0 and Col-0 35S::*Pb*$GH_3$ lines at 3 dpi (scale bars = 50 µm).

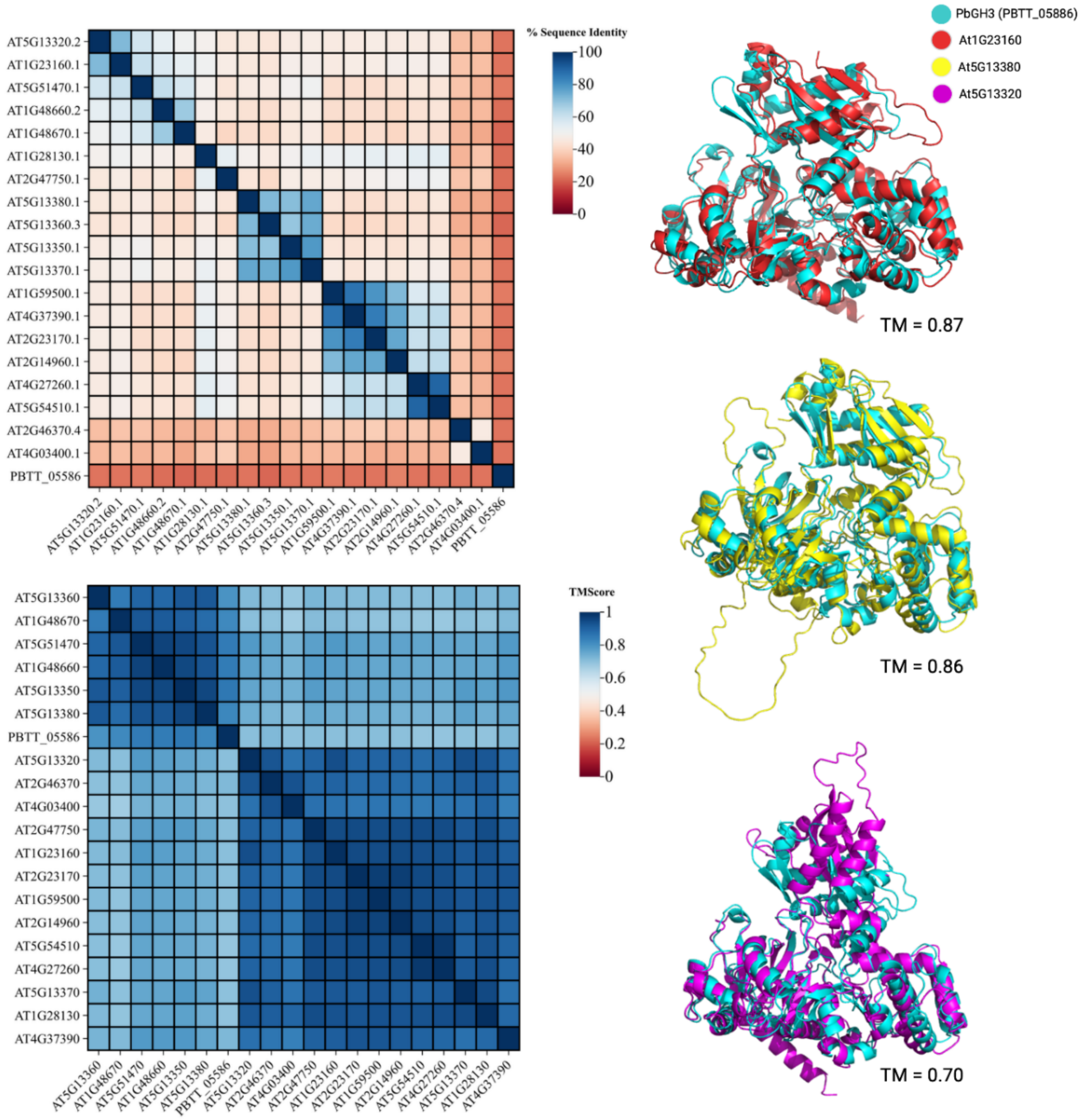


**Figure S10.** Sequence and structural conservation of *Pb*GH$_3$ relative to *Arabidopsis* GH$_3$ family members. (**A**) Heatmap showing pairwise percentage of amino-acid sequence identity between *Arabidopsis thaliana* GH$_3$ proteins and the *Plasmodiophora brassicae* effector *Pb*GH$_3$ (PBTT_05586). Despite overall low primary sequence identity, *Pb*GH$_3$ shows detectable similarity to multiple GH$_3$ family members. (**B**) Heatmap of TM-scores derived from pairwise structural alignments, indicating high three-dimensional similarity between *Pb*GH$_3$ and plant GH$_3$ proteins (TM-score > 0.7), consistent with a conserved protein fold. (**C–E**) Structural superpositions of *Pb*GH$_3$ (cyan) with representative *Arabidopsis* GH$_3$ proteins: *At*GH$_{3.1}$ (AT1G23160) (red), *At*GH$_{3.8}$ (AT5G13380) (yellow), and *At*GH$_{3.2}$ (AT5G13320) (magenta). TM-scores are indicated for each alignment, revealing strong structural conservation despite low sequence identity.

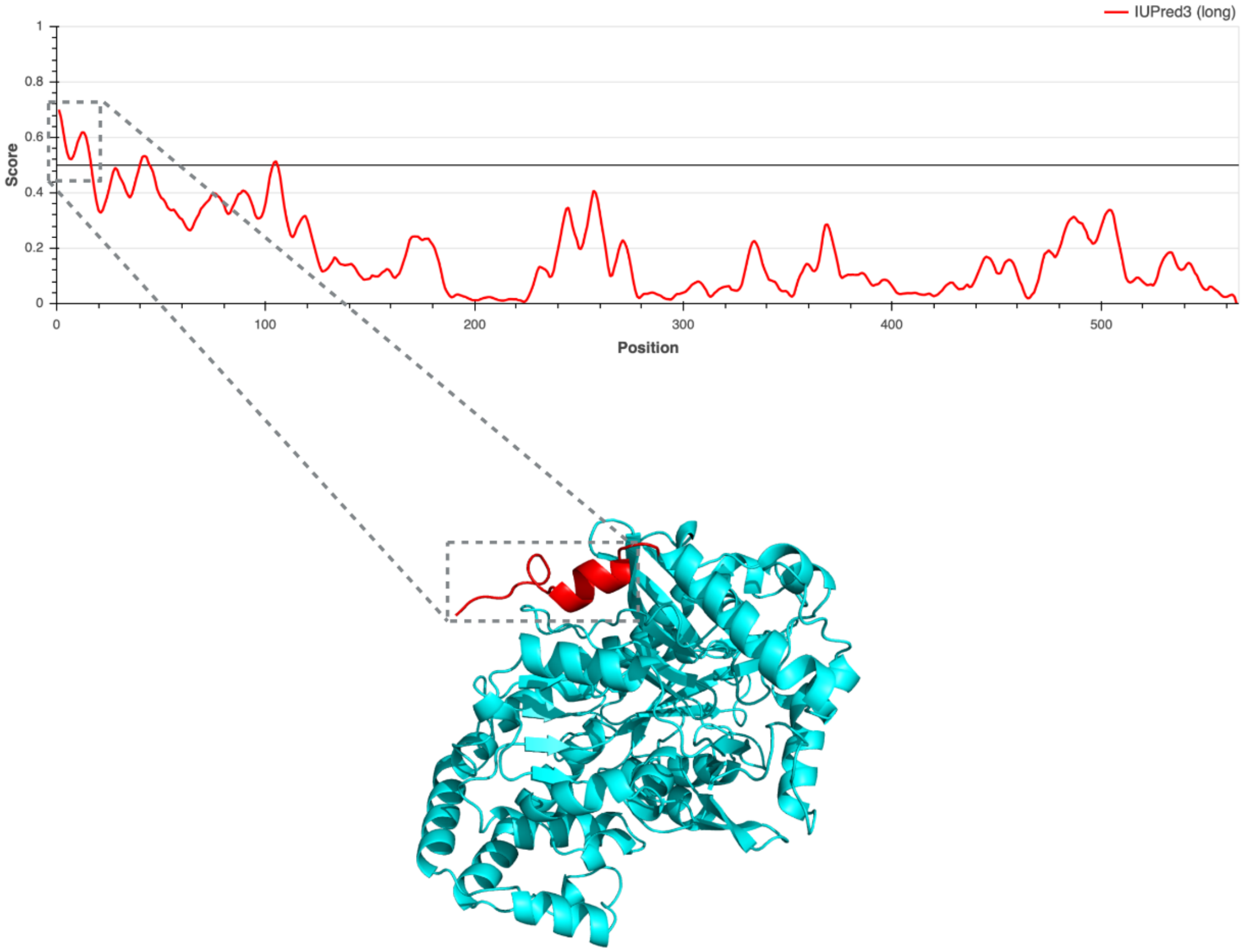


**FIGURE S11.** Predicted disordered N-terminal region of *Pb*$GH_3$. The red curve indicates the disorder score for each amino acid position, with values above the threshold (grey line) indicating predicted disordered regions. The first ~16 amino acids of *Pb*$GH_3$ display a high disorder score, suggesting the presence of a flexible N-terminal region. The corresponding region is highlighted in red on the predicted *Pb*$GH_3$ protein structure, indicating a potentially unstructured segment that may contribute to protein interactions, targeting, or regulatory functions.
